# Conflicts and coalitions among selfish toxin-antidote elements shape the evolution of hyperdivergent haplotypes

**DOI:** 10.64898/2026.10.22.760680

**Authors:** Alevtina Koreshova, Andreas Hagmüller, Arda Kara, Pinelopi Pliota, Tianhua Liao, George Kobakhidze, Joseph Gokcezade, Peter Duchek, Christian Braendle, Gang Dong, Alejandro Burga

## Abstract

Genetic conflict drives evolutionary innovation, yet most research has focused on arms races between selfish elements and their hosts. A second layer—competition among selfish elements—remains largely unexplored. Here, we uncover such a conflict between two antagonistic toxin–antidote elements (TAs), *loki-1/sgyn-1* and *thor-1/sif-1*, in *Caenorhabditis tropicalis*. Each toxin, LOKI-1 or THOR-1, selectively kills progeny homozygous for the opposing haplotype, favoring heterozygotes. Their antidotes, SGYN-1 and SIF-1, evolved from F-box proteins, although SIF-1 subsequently lost its F-box domain. We solved the crystal structure of the THOR-1–SIF-1 complex and used *in vivo* mutagenesis to validate the binding interface, supporting direct toxin neutralization independently of F-box-mediated degradation. A survey of 676 wild isolates revealed two contrasting evolutionary outcomes of antagonizing TAs: independently evolved *loki-1/sgyn-1* and *thor-1/sif-1* remain confined to alternative hyperdivergent haplotypes, whereas other homologous TAs can recombine to form composite haplotypes. Altogether, our findings establish competition among selfish elements as a distinct form of genetic conflict, and support models in which antagonistic TAs act as barriers that restrict gene flow between neighboring populations.

## Main text

The molecular arms race between selfish elements and their host has been central to our understanding of how these genetic parasites shape genome evolution and drive biological innovation^1^. Yet what persists together is not necessarily shaped by common interests. While useful, this framework misses an important dimension of their biology. Within every genome, multiple selfish elements coexist and may compete—or even antagonize one another—creating dynamics whose ecological and evolutionary consequences remain largely unexplored. A major limiting factor is the genetic and molecular identification of selfish elements engaged in such intra-parasitic conflicts. Here, we address this gap by dissecting and characterizing a pair of antagonistic toxin–antidote elements in the nematode *Caenorhabditis tropicalis*.

Toxin–antidote elements (TAs) are selfish genes that increase their frequency in populations by selectively poisoning individuals that do not inherit them^2^. In nematodes, TAs typically consist of two genes in tight genetic linkage: a maternally expressed toxin and a zygotically expressed antidote. In crosses between carrier and non-carrier strains, F_1_ heterozygous mothers are unaffected, but they poison all of their progeny by loading the toxin into unfertilized eggs. Only individuals that inherit at least one copy of the element can express the antidote and neutralize the toxin. As a result, homozygous non-carrier individuals in the F_2_ generation are poisoned, with phenotypes ranging from developmental delay to embryonic lethality. Over successive generations, this gene drive leads to the spread and eventual fixation of the TA in the population. To date, three TAs have been identified in *C. elegans*: *peel-1/zeel-1* (Chr. I), *sup-35/pha-1* (Chr. III), and *mll-1/smll-1* (Chr. V)^3–5^. While these elements efficiently distort inheritance, they occupy different chromosomes and therefore spread independently of one another. In contrast, recent work in *C. tropicalis*, a relative of *C. elegans*, has revealed a more complex landscape in which TAs segregate as alternative haplotypes at homologous genomic loci or are located in close proximity to each other, creating opportunities for conflict between selfish elements with opposing transmission interests^6,7^.

Crosses between two Caribbean *C. tropicalis* isolates, NIC203 (Guadeloupe) and EG6180 (Puerto Rico), revealed a high density of selfish elements, with at least seven active TAs segregating across just three chromosomes. Five have been characterized at the molecular level, including the homologous TAs *slow-1/grow-1* and *slow-2/grow-2* on Chr. III of NIC203 and EG6180, respectively^8,9^. Yet two additional TAs in EG6180, located on Chr. II and V, remained molecularly unresolved. To fill this gap, we set out to dissect the TA on the right arm of Chr. V. This effort led to the identification of *loki-1/sgyn-1* and, unexpectedly, a previously unrecognized TA in NIC203, *thor-1/sif-1*. The newly identified TAs, *loki-1/sgyn-1* and *thor-1/sif-1* are not homologous and reside within non-recombining hyperdivergent haplotypes, placing them in direct evolutionary conflict. Although both antidotes, SGYN-1 and SIF-1, evolved from F-box proteins, SIF-1 lost its F-box domain. To investigate the molecular basis of SIF-1 antidote activity, we solved the crystal structure of the THOR-1–SIF-1 complex, revealing the interface that mediates toxin neutralization. We further identified shared molecular and regulatory features and surveyed the distribution and recombination patterns of antagonistic TAs across 676 wild isolates worldwide, revealing two distinct evolutionary strategies that shape TA diversity in natural populations.

### A novel pair of antagonistic TAs in *C. tropicalis*

To map selfish TAs present in the EG6180 wild-isolate background, we previously employed a multigenerational backcross strategy^6^. Briefly, EG6180 hermaphrodites were crossed with NIC203 males, and the resulting F_1_ hybrid hermaphrodites were repeatedly backcrossed to NIC203 males for more than 30 generations (Figure 1A). This strategy progressively replaced most of the genome with NIC203 alleles, except for EG6180 maternal-effect TA loci, which were retained because they selectively poison progeny homozygous for the NIC203 non-carrier susceptible haplotype. This mating scheme resulted in a nearly isogenic line (NIL) carrying two putative TAs, one in Chr. II and a second in Chr. V. To facilitate their characterization, we derived NILs carrying each introgression separately. Because both loci act through maternal-effect killing, we used a crossing scheme that avoided toxin-mediated lethality during NIL derivation by crossing NIC203 hermaphrodites with males heterozygous for both introgressions. In this cross, maternally expressed toxins cannot eliminate homozygous non-carrier progeny, allowing recovery of the desired genotypes. Consistent with this expectation, we readily derived a homozygous Chr. II NIL. In contrast, attempts to derive a Chr. V NIL revealed an unexpected pattern of inheritance. Rather than exhibiting the segregation distortion expected for a single maternal-effect TA, progeny derived from heterozygous mothers displayed a striking excess of heterozygous progeny, suggesting that both homozygous classes were being selectively eliminated (Figure 1A). Nevertheless, we recovered a rare homozygous Chr. V NIL that was fully viable and fertile, indicating that homozygosity for the introgressed region itself was not deleterious (96,5% wild-type, n=200; Figure 1B).

**Figure 1.**
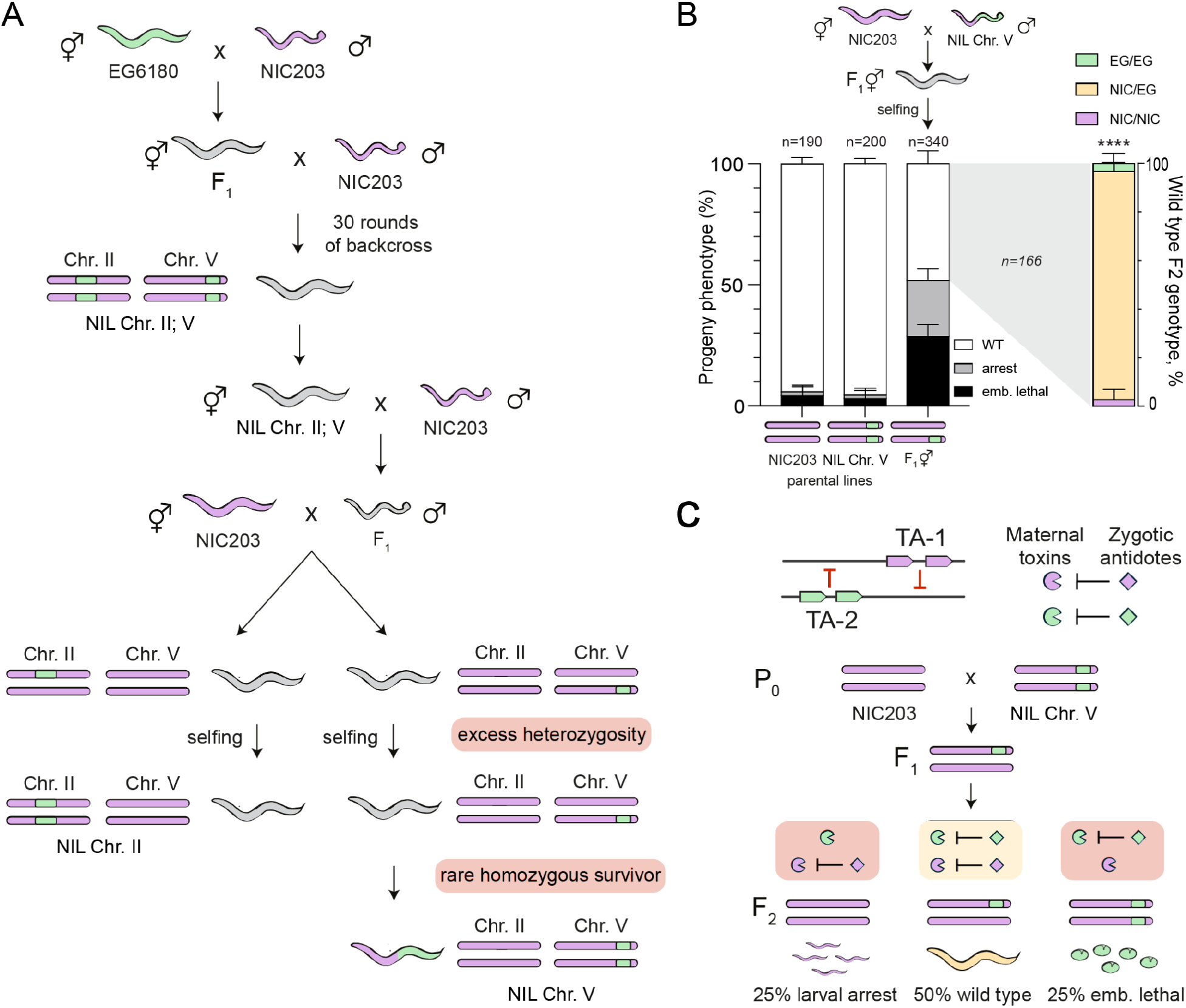
A novel pair of antagonistic TAs in C. tropicalis Chr. V. **(A)** Schematic of the crossing strategy used to generate Near-Isogenic Lines (NILs) carrying either a Chr. II or Chr. V EG6180 TA element. **(B)** In crosses between NIC203 hermaphrodites and NIL Chr. V males approximately half of their F_2_ progeny died before reaching adulthood (∼51%; n=340). Dead progeny corresponded to two distinct phenotypic classes: embryonic lethal (black) and larval lethal (grey). Genotyping of the introgressed region revealed a large excess of heterozygous individuals among the F_2_ wild-type progeny (P < 0.0001, exact multinomial goodness-of-fit test to the expected Mendelian ratio). For this and all following figures: \*\*\*\**P < 0.0001*, ns = not significant. **(C)** Proposed genetic model for the inheritance pattern observed in (B) consisting of a pair of antagonistic TAs. In crosses between two strains carrying unique TAs, all F_2_ individuals are maternally-poisoned with two toxins. Only heterozygous F_2_ individuals express both zygotic antidotes and are not affected (wild-type phenotype). In contrast, homozygous individuals can only neutralize one toxin, leading to 50% affected F_2_ progeny.

To determine the basis of this distortion, we performed diagnostic crosses using the Chr. V NIL. When NIC203 hermaphrodites were crossed to Chr. V NIL males, we observed ∼50% of lethality among their F_2_ progeny. Further, two distinct classes of lethality were observed: approximately half of the affected progeny arrested and died during the L1/L2 larval stages, whereas the remainder died during embryogenesis (28,5% embryonic lethality, 22,6% larval arrest; n=340; Figure 1B). Genotyping of the surviving F_2_ progeny revealed that nearly all wild-type progeny were heterozygous for the NIL introgression (NIC/EG) (Figure 1B). Thus, in addition to the expected elimination of homozygous non-carriers (NIC/NIC) by the EG6180 TA, homozygous carriers of the introgression (EG/EG) were also selectively eliminated. The loss of both homozygous classes resulted in a strong excess of heterozygotes, consistent with the segregation distortion observed during derivation of the parental Chr. V NIL (Figure 1A).

The selective depletion of EG/EG individuals strongly suggested the presence of a second active TA (Figure 1C). Because this second activity specifically targeted progeny homozygous for the EG6180 introgression, we predicted that this element should reside in the NIC203 genome, within the Chr. V region homologous to the EG6180 introgression. In this model, two distinct TA elements carried by EG6180 and NIC203 occupy homologous Chr V haplotypes and act in direct antagonism. Heterozygous F_1_ mothers deposit both toxins into all F_2_ eggs, however only heterozygous F_2_ progeny inherit and express both zygotic antidotes, allowing them to survive (Figure 1C). To test this hypothesis, we set out to fine-map the underlying loci.

### Genetic mapping of the *loki-1/sgyn-1* TA

The Chr. V NIL carries a large EG6180-derived introgression spanning ∼1.5 Mb and encompassing over 300 genes. To narrow the list of candidate genes, we crossed Chr. V NIL hermaphrodites to NIC203 males and screened for recombination events within the introgressed region. From this screen, we recovered four recombinant lines carrying smaller, overlapping segments of the original introgression (Figure 2A; see Methods). Each of these four sub-NILs was then crossed to the original Chr. V NIL, and embryonic and larval lethality among the resulting progeny was scored as a readout of TA activity.

**Figure 2.**
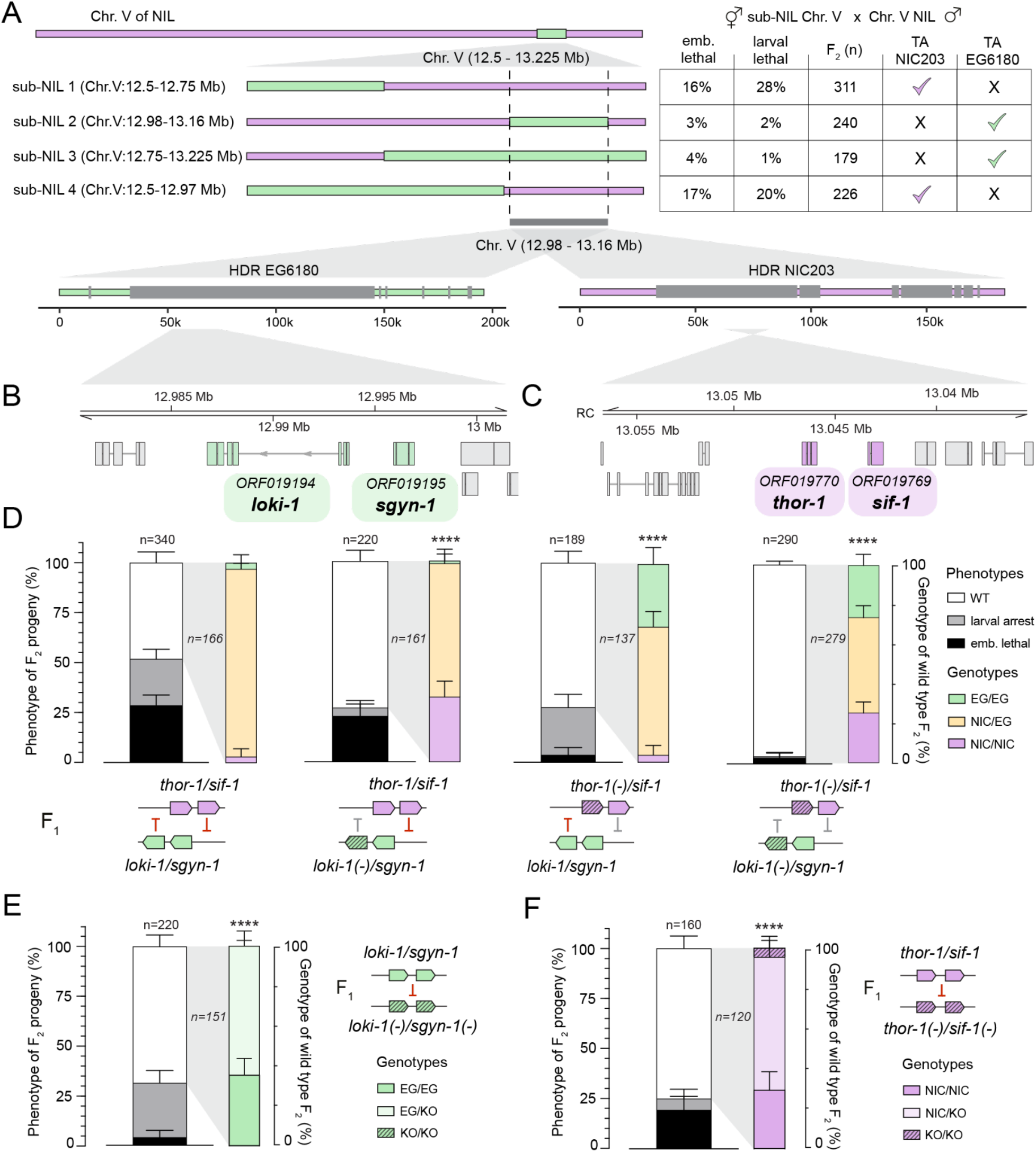
Genetic mapping of the *loki-1/sgyn-1* and *thor-1/sif-1* antagonistic TAs. **(A)** Schematic of the recombinant lines used to map the antagonistic TAs on Chr. V. Starting from the parental Chr. V NIL, four recombinant sub-NILs were generated and crossed with Chr. V NIL males. The activity of the EG6180 or NIC203 TA was inferred from the presence of dead larvae or embryos, respectively, among the F_2_ progeny. Gray boxes indicate the EG6180 and NIC203 hyperdivergent regions (HDRs) within the mapped interval (bottom). **(B)** Enlarged view of the region containing the candidate genes for the EG6180 TA: *ORF019194* and *ORF019195*, corresponding to the toxin *loki-1* and antidote *sgyn-1*, respectively. Bottom track shows aligned reads from EG6180 mRNA-seq. **(C)** Enlarged view of the region containing the candidate genes for the NIC203 TA: *ORF019770* and *ORF019769*, corresponding to the toxin *thor-1* and antidote *sif-1*, respectively. RC denotes reverse-complement orientation. **(D)** Genetic crosses used to identify the toxins encoded by the EG6180 and NIC203 TA elements. From left to right: Chr. V NIL hermaphrodites crossed with EG6180 males; Chr. V NIL hermaphrodites crossed with *loki-1(-)* EG6180 males; *thor-1(-)* Chr. V NIL hermaphrodites crossed with EG6180 males; and *thor-1(-)* Chr. V NIL hermaphrodites crossed with *loki-1(-)* EG6180 males. F_2_ phenotypes were quantified for each cross, and only phenotypically wild-type F_2_ progeny were genotyped. *P* values calculated using exact multinomial goodness-of-fit test to the cross between NIC203 hermaphrodites and Chr. V NIL males. The wild-type F_2_ genotype distribution in the cross between *thor-1(-)* Chr. V NIL hermaphrodites and *loki-1(-)* EG6180 males was not significantly different from Mendelian expectation (*P = 0.79*). **(E)** Cross between wild type EG6180 hermaphrodites and double mutant *loki-1(-);sgyn-1(-)* EG6180 males. F_2_ phenotypes were quantified for each cross, and only phenotypically wild-type F_2_ progeny were genotyped. *P* values calculated using exact multinomial goodness-of-fit test to the expected Mendelian ratio. **(F)** Cross between wild type NIC203 hermaphrodites and double mutant *thor-1(-);sif-1(-)* NIC203 males. F_2_ phenotypes were quantified for each cross, and only phenotypically wild-type F_2_ progeny were genotyped.

Comparison of the resulting inheritance patterns localized both elements to a shared ∼180-kb interval on Chr. V (12,98-13,16 Mb) (Figure 2A).

To identify candidate toxin and antidote genes, we examined pairs of genes missing or mutated in the susceptible haplotype that also exhibit signatures of rapid evolution (see Methods). Among the strongest candidates within the EG6180 mapped interval, we identified the adjacent genes *ORF019194* and *ORF019195* (see methods; Table S1). *ORF019194* lacks detectable homology to known genes, whereas *ORF019195* encodes a predicted F-box protein, like other previously characterized antidotes^9^. To test whether *ORF019194* codes for the toxin, we generated a null allele by CRISPR/Cas9-mediated homologous repair in the Chr. V NIL background. The *ORF019194* mutants were healthy and did not show any abnormal phenotype (94,5% wild-type; n=200; Figure S1). When NIC203 hermaphrodites were crossed to *ORF019194* NIL mutant males, lethality among homozygous non-carriers (NIC/NIC) was completely abolished in the F_2_ generation and larval arrest was no longer observed (22,3% embryonic lethality, 4,5% larval arrest; n=220; Figure 2D). In contrast, homozygous carriers of the introgression (EG/EG) continued to be selectively eliminated: 80% (16/20) of genotyped dead embryos were EG/EG, consistent with the hypothesis that the opposing NIC203-derived TA acts independently (Figure 2D). These results demonstrate that *ORF019194* is required for the toxic activity of the EG6180 TA element and therefore corresponds to the toxin.

We next tested whether *ORF019195* codes for the antidote. To do so, we generated a *ORF019195* null mutation in the *ORF019194* toxin mutant background. The double-mutant line was healthy and showed only background levels of embryonic lethality (94,6% wild-type; n=130; Figure S1). We then crossed double-mutant Chr. V NIL hermaphrodites to wild-type Chr. V NIL males, and analyzed the resulting F_2_ progeny. If *ORF019195* functions as the antidote, the double-mutant allele should behave as a susceptible haplotype and therefore be selectively eliminated by the toxin inherited from the wild-type NIL parent^6^. This is precisely what we observed: progeny homozygous for the double-mutant haplotype were selectively lost and ∼25% of the F_2_ progeny died as larvae (27,3% larval arrest; n=220; Figure 2E). Together, these experiments demonstrate that *ORF019194* encodes a toxin, which we named *loki-1* (for ***L****arval stage **O**ffspring **KI**ller*) and *ORF019195* encodes its cognate antidote, which we named *sgyn-1* (for ***S****uppressor of **G**enetic incompatibilit**Y** with F-box domai**N***, pronounced *SIG-in*).

### Genetic mapping of the *thor-1/sif-1* TA

Having mapped the *loki-1/sgyn-1* TA, we set out to identify the putative antagonistic TA in NIC203. Applying the same candidate prioritization strategy, we identified the adjacent genes *ORF019770* and *ORF019769* as the strongest candidates for the NIC203 toxin and antidote, respectively (see methods; Table S2). To test whether *ORF019770* encodes the toxin, we generated an *ORF019770* null allele using CRISPR/Cas in the NIC203 background. The *ORF019770* mutants were healthy and did not show any abnormal phenotype (94,5% wild-type; n=203; Figure S1). We then crossed mutant hermaphrodites to Chr. V NIL males. Loss of *ORF019770* abrogated embryonic lethality in the F_2_ generation (3,7% embryonic lethality, 23,8% larval arrest; n=189; Figure 2D), and homozygous carriers of the EG6180 introgression (EG/EG) were recovered at the expected Mendelian frequency among viable progeny, demonstrating that *ORF019770* is the toxin. In contrast, homozygous carriers of the NIC203 haplotype continued to be selectively eliminated: 96% (23/24) of genotyped dead larvae were NIC/NIC homozygotes.

Next tested whether *ORF019769* functions as the antidote. To do so, we generated an *ORF019770; ORF019769* double-mutant in the NIC203 background and crossed wild-type NIC203 hermaphrodites to double-mutant NIC203 males (Figure 2F). We found that progeny homozygous for the double-mutant haplotype were selectively eliminated, resulting in approximately 25% embryonic lethality among the F_2_ progeny (19,4% embryonic lethality; n=160; Figure 2E). In contrast, the parental double-mutant line was healthy and showed only background levels of embryonic lethality (98,8% wild-type; n=170; Figure S1). Together, these experiments established *ORF019770* and *ORF019769* as a novel NIC203 TA pair, with *ORF019770* encoding the toxin and *ORF019769* its cognate antidote. Accordingly, we renamed *ORF019770* and *ORF019769* as *thor-1* (for ***T****oxic **H**ybrid **O**ffspring **R**atio distorter*) and *sif-1* (for ***S****uppressor of genetic **I**ncompatibility without **F**-box domain*), respectively.

Next, we asked whether the gene drive activity of *loki-1/sgyn-1* and *thor-1/sif-1* fully accounts for the large excess of heterozygous individuals among the F_2_ progeny of the cross between NIC203 hermaphrodites and Chr. V NIL males. To do so, we crossed *thor-1(-)* mutant NIC203 hermaphrodites to *loki-1(-)* mutant Chr. V NIL males and inspected the resulting F_2_ progeny. In the absence of both toxins, we observed only background levels of lethality (3,1% embryonic lethality, 0,7% larval arrest; n=290; Figure 2D), and critically, heterozygotes and both homozygous classes (NIC/NIC and EG/EG) were recovered at the expected Mendelian frequencies (25,2% NIC/NIC, 48,1% NIC/EG, 26,7% EG/EG; n=262), indicating that the apparent heterozygote advantage at this locus stemmed from the antagonistic activity of the two selfish TAs.

### The *thor-1* toxin is epigenetically licensed

The *loki-1/sgyn-1* TA is equally active regardless of the sex of the carrier parent, as crosses with either Chr. V NIL hermaphrodites or males result in poisoning of the non-carrier F_2_ (Figure 3A). In contrast, the *thor-1/sif-1* TA appears to be only active when the TA is inherited through the maternal lineage, indicative of a parent-of-origin effect (Figure 3A). This inheritance pattern closely resembles that of the *slow-1/grow-1* TA on Chr. III^8^. In the *slow-1/grow-1* system, paternal inheritance of the TA leads to epigenetic repression of the *slow-1* toxin by the piRNA pathway in the germline of heterozygous F_1_ mothers. Consequently, maternal deposition of SLOW-1 is reduced and their progeny is not poisoned even in the absence of the antidote. By contrast, when the TA is maternally inherited, maternal *slow-1* transcripts deposited in the oocyte protect the zygotic copy from piRNA-mediated silencing through a process termed epigenetic licensing^8,10^.

**Figure 3.**
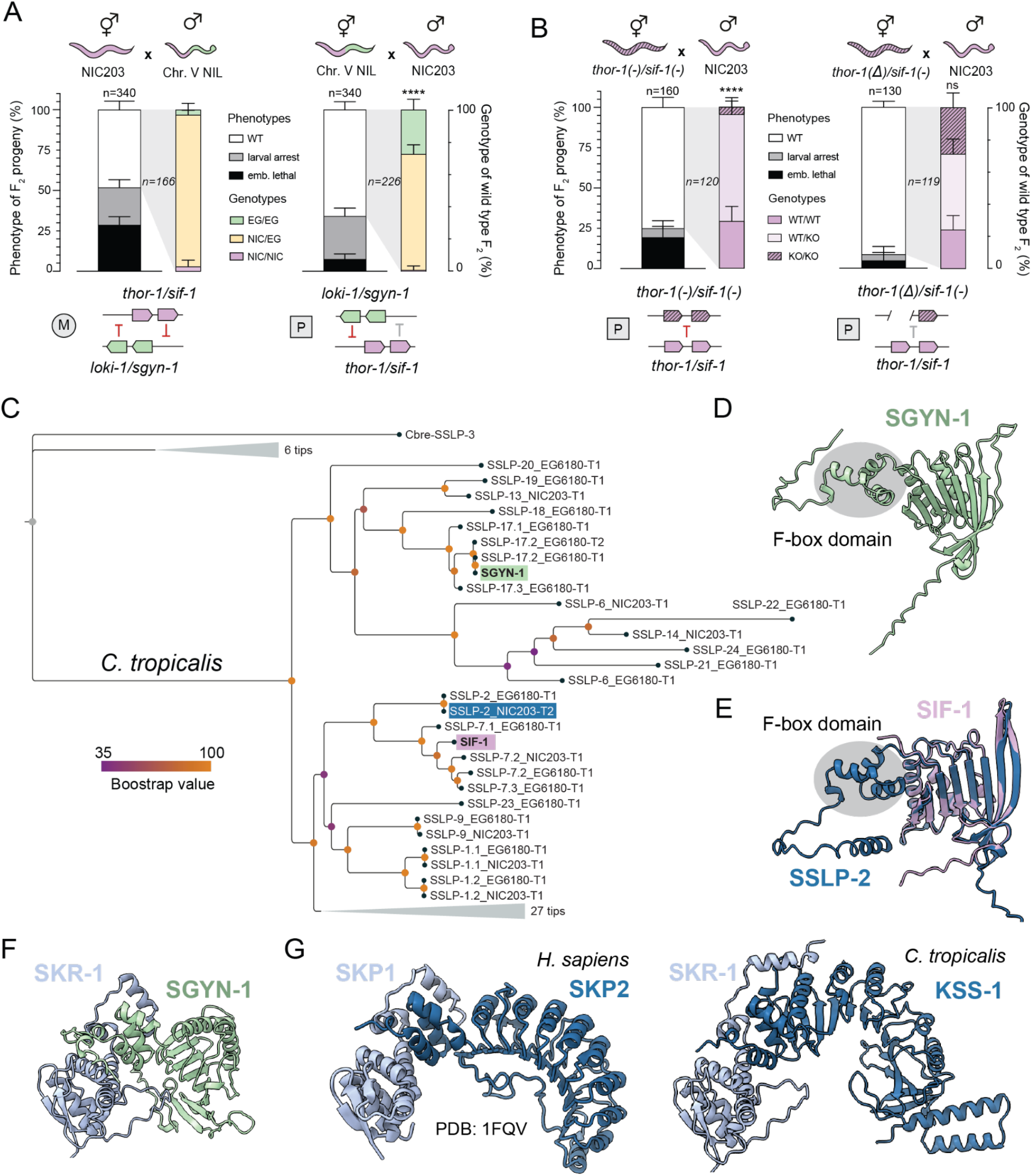
Epigenetic licensing of the *thor-1* toxin and evolution of SGYN-1 and SIF-1 from a rapidly diverging F-box protein family. **(A)** Crosses between NIC203 hermaphrodites and Chr V NIL males (right), and the reciprocal crosses (left), reveal a parent-of-origin effect on *thor-1* toxicity. Maternal (M) or paternal (P) inheritance refers to the *thor-1/sif-1* locus. *P* values calculated using exact multinomial goodness-of-fit test to the cross between NIC203 hermaphrodites and Chr. V NIL males. **(B)** Crosses between *thor-1(-); sif-1(-)* (left) or *thor-1(Δ); sif-1(-)* (left) double-mutant NIC203 hermaphrodites and wild-type NIC203 males. The *thor-1(−)* allele contains a frameshift mutation, whereas *thor-1(Δ)* is a deletion of the entire *thor-1* gene.Paternal (P) inheritance refers to the wild-type *thor-1/sif-1* NIC203 locus. *P* values calculated using exact multinomial goodness-of-fit test to the expected Mendelian ratio. **(C)** Phylogenetic relationships among the SGYN-1 and SIF-1 antidotes and the *C. tropicalis* SSLP family of intact F-box proteins and proteins that lost their F-box domains like SIF-1. SSLP proteins from *C. brenneri* were included as an outgroup. **(D)** AlphaFold 3-predicted structure of SGYN-1 (light green), with its N-terminal F-box domain highlighted in gray. **(E)** Structural alignment of the AlphaFold 3-predicted models of SIF-1 (lilac) and the closely related SSLP-2 protein (blue). The N-terminal F-box domain of SSLP-2, which is absent from SIF-1, is highlighted in gray. **(F)** AlphaFold 3-predicted interaction between SKR-1 (residues 1-10 not shown) and SGYN-1, mediated by the predicted F-box domain of SGYN-1. **(G)** X-ray crystal structure of human SKP1 bound to the SKP2 F-box protein (PDB: 1FQV; left) and AlphaFold 3-predicted structure of *C. tropicalis* SKR-1 bound to the KSS-1 antidote (right).

To determine whether *thor-1* is similarly subject to epigenetic licensing, we compared the effects of a *thor-1(-)* null frameshift allele and a *thor-1(Δ)* deletion allele that removes the entire *thor-1* locus (5.8 kb) (Figure 3B). As epigenetic licensing depends on the presence of maternal mRNA rather than its translation product, a frameshift allele retains licensing activity despite abolishing THOR-1 function, whereas a complete deletion should eliminate licensing by removing all maternal *thor-1* transcripts^8^. We therefore crossed either *thor-1(-); sif-1(-)* or *thor-1(Δ); sif-1(-)* double-mutant hermaphrodites to NIC203 males carrying the wild-type *thor-1/sif-1* TA and assayed the activity of the paternally inherited TA. When mothers carried the *thor-1(-)* frameshift allele, the paternally inherited *thor-1/sif-1* TA was fully active, indicating that maternal *thor-1* mRNA but not THOR-1 protein, was sufficient to license the zygotic toxin (Figure 3B). In contrast, when mothers carried the *thor-1(Δ)* allele, the paternally inherited TA was inactive (*P = 0.83*; Figure 3B). This was evidenced by the absence of embryonic lethality and the recovery of F_2_ progeny homozygous for the susceptible *thor-1(Δ); sif-1(-)* genotype at the expected Mendelian frequency (Figure 3B). Together, these results indicate that zygotic *thor-1*, like *slow-1*, requires maternally supplied transcripts to prevent its epigenetic repression following paternal inheritance of the TA, consistent with regulation by epigenetic licensing. Further, this mechanism explains the sudden activation of the *thor-1/sif-1* TA while deriving the Chr. V NIL. During the backcross procedure, we employed NIC203 males, and thus, *thor-1/sif-1* TA was epigenetically repressed (Figure 1A). However, when deriving the Chr. II and Chr. V NILs, we employed NIC203 hermaphrodites, resulting in *thor-1* licensing and an active *thor-1/sif-1* TA (Figure 1A).

### The SGYN-1 and SIF-1 antidotes evolved from F-box proteins

Despite their close proximity on Chr. V, *loki-1/sgyn-1* and *thor-1/sif-1* do not appear to share a common evolutionary origin. Their respective toxins, LOKI-1 and THOR-1, show no significant sequence similarity to one another. However, both proteins show hallmarks of rapid evolution in line with their selfish role. *loki-1* is a chimeric transcript; its first three exons do not contribute to the protein due to the absence of start codons and presence of multiple stop codons. Further, they are separated from the other downstream exons by a 5 kb intron. A BLASTN search didn’t yield any significant hits within the EG6180 genome, so we performed a BLASTP search with the best predicted protein sequence. The best hit from this search (e-value = 3×10^−30^) was a predicted checkpoint kinase homologous to *chk-2* in *C. elegans* and Loki in *Drosophila melanogaster*, located ∼650 kb away from *loki-1*. The remaining four exons of *loki-1* encode a 309-amino-acid protein that lacks any recognizable protein domains. A BLAST search against the *C. tropicalis*proteome identified a single significant hit (e-value = 9×10^−16^): the gene immediately downstream to *loki-1*. This gene, *ORF019193*, is homologous to *C. elegans ucr-2.3*, which encodes a predicted mitochondrial processing peptidase (Figure S2). However, LOKI-1 itself is highly divergent from ORF019193, and AlphaFold3 failed to generate a high-confidence structural model of this toxin, consistent with extensive sequence divergence (Figure S2). THOR-1 is an even more enigmatic protein. At only 134 amino acids in length, it shows no detectable sequence similarity to any other protein in *C. tropicalis* or in any other publicly available animal, plant, or bacterial genomes. Its extreme lack of homology suggests that THOR-1 may represent a *de novo*–evolved protein or, alternatively, may have been acquired through horizontal gene transfer^11^. However, even the latter scenario would require substantial sequence divergence following acquisition to explain the absence of detectable homologs.

The antidotes, SGYN-1 and SIF-1, also show signatures of rapid evolution. But unlike the toxins, they belong to a well-characterized protein family: F-box proteins. F-box proteins function as substrate-recognition modules of the SCF (Skp1–Cullin-1–F-box) E3 ubiquitin ligase complex, directing specific proteins for ubiquitination and subsequent degradation. This role relies on two key interactions: binding the Skp1 adaptor through a conserved N-terminal F-box domain and recognizing target substrates through a specialized and variable protein–protein interaction domain^12–14^. In this way, F-box proteins bridge the SCF complex to its substrates, conferring specificity to ubiquitin-mediated protein degradation. In nematodes, F-box proteins have undergone remarkable lineage-specific expansions, with individual species often encoding hundreds to thousands of family members^15,16^. In addition, they evolve at an exceptionally rapid pace, mainly through tandem duplications, resulting in extensive sequence diversification and limited conservation even between closely related species. Despite sharing limited sequence identity, a systematic search for SGYN-1 and SIF-1 homologous across *Caenorhabditis* species revealed that both antidotes belong to the same family of F-box proteins (Figure 3C and S3). This family, which we named *sslp* (for Sif and Sgyn-Like Proteins), is relatively small and only present in *C. tropicalis* and *C. brenneri* (Table S3). SGYN-1 has a conserved F-box domain on its N-terminus and Alphafold3 predicts high-confidence interactions with Skp1 homologs, with *C. tropicalis* SKR-1 as its strongest predicted binding partner (Figure 3D, Figure 3F, and Table S4). These findings suggest that SGYN-1 may function as the substrate-recognition component of an SCF E3 ubiquitin ligase complex. This proposed neutralization mechanism resembles that of the KSS-1 antidote, which binds both SKR-1 and its cognate toxin, KLMT-1, targeting the latter for degradation^9^. Despite lacking detectable homology, SGYN-1 and KSS-1 also exhibit structural similarities to the human F-box protein SKP2: all three possess a concave surface formed predominantly by β-strands, although this surface is considerably smaller in SGYN-1 (Figure 3G). SIF-1, in contrast, exhibits a striking departure from the canonical F-box architecture. Comparison with closely related F-box proteins in *C. tropicalis* revealed that SIF-1 has completely lost the N-terminal F-box domain (Figure 3E). It is therefore unlikely to bind Skp1 or recruit the SCF E3 ubiquitin ligase complex. Thus, although SIF-1 evolved from an F-box protein, its loss of the F-box domain suggests that it has been repurposed to neutralize THOR-1 through a mechanism independent of ubiquitin-mediated degradation.

### Structural characterization of the THOR-1–SIF-1 neutralization complex

To investigate the molecular basis of SIF-1 antidote activity, we co-expressed untagged THOR-1 together with N-terminally tagged 6xHis::SIF-1 in Sf9 insect cells (Figure 4A). Following affinity purification using a HisTrap column and subsequent size-exclusion chromatography, analysis of the eluate by SDS–PAGE revealed two major bands: a ∼25 kDa band corresponding to the tagged antidote and a ∼15 kDa band corresponding to the toxin (Figure 4A). The co-purification and relative abundance of the untagged toxin strongly suggested that SIF-1 physically interacts with THOR-1 and forms a stable complex (Figure 4A). To further characterize this complex, we performed size-exclusion chromatography coupled to multi-angle light scattering (SEC–MALS). The purified THOR-1–SIF-1 complex had a molecular mass of 39.6 ± 0.17 kDa (n = 3), consistent with the theoretical 38.8 kDa expected for a 1:1 heterodimer (Figure 4B). This stoichiometry was further supported by the single major SEC peak containing both THOR-1 and SIF-1 (Figure 4A).

**Figure 4.**
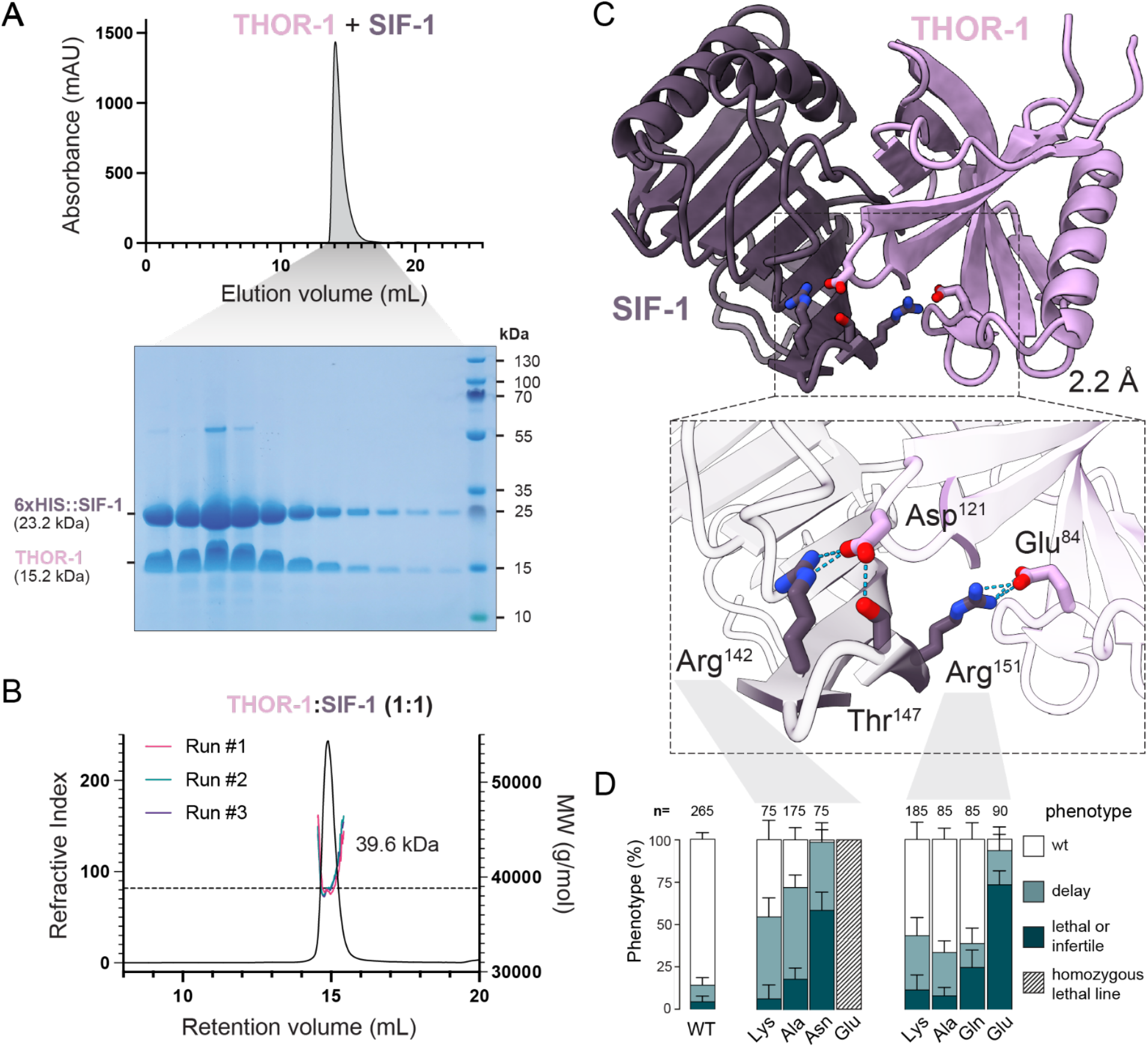
Biochemical characterization and X-ray structure of the THOR-1–SIF-1 neutralization complex. **(A)** Top, gel filtration of the THOR-1–6×His::SIF-1 complex expressed in Sf9 cells following HisTrap affinity purification and ion-exchange chromatography . Bottom, SDS–PAGE and Coomassie staining of the indicated elution fractions. **(B)** Size-exclusion chromatography coupled with multi-angle light scattering (SEC–MALS) analysis of the THOR-1–SIF-1 complex. The measured molecular mass was 39.6 ± 0.17 kDa (mean ± [SD/SEM], n = 3). **(C)** X-ray crystal structure of the THOR-1 toxin bound to the SIF-1 antidote. The inset shows a magnified view of the electrostatic interaction interface. SIF-1 Arg^142^ forms a salt bridge with THOR-1 Asp^121^, while SIF-1 Thr^147^ forms a hydrogen bond with the same residue. SIF-1 Arg^151^ forms an additional salt bridge with THOR-1 Glu^84^. **(D)** Phenotypes of *sif-1* mutant lines carrying substitutions at the toxin-interacting residues Arg^142^ and Arg^151^. *n* indicates the total number of individuals phenotyped for each mutant line.

Next, we sought to determine the structure of the THOR-1–SIF-1 complex. Initial crystallization trials were unsuccessful. Inspection of predicted structures revealed flexible terminal extensions in both proteins (Figure S4A), suggesting that conformational heterogeneity might hinder crystallization. We therefore performed limited proteolysis with several proteases to identify conditions that removed flexible regions while preserving the structured cores of the complex. All tested proteases generated discrete, protease-resistant products, with thermolysin producing particularly well-defined and stable fragments (see Methods and Figure S4B). Thus, we incorporated thermolysin-mediated in situ proteolysis into subsequent crystallization trials, which yielded well-diffracting crystals (Figure S5). The structure of the THOR-1–SIF-1 complex was subsequently determined at 2.2 Å resolution (Table S5).

The refined crystal structure contains residues 18-133 of THOR-1 and residues 6-175 of SIF-1 and reveals an extensive protein–protein interface between the toxin and antidote (Figure 4C; Figure S6A). A prominent feature is an intermolecular β-sheet formed between SIF-1 residues 100–102 and THOR-1 residues 32–36 (Figure S6B). In addition, multiple hydrogen bonds and salt bridges contribute to stabilizing the interface (Figure S6A). Among these interactions, SIF-1 Arg^142^ and Thr^147^ bind to THOR-1 Asp^121^ via a salt bridge and hydrogen bond, whereas Arg^151^ forms a salt bridge with THOR-1 Glu^84^ (Figure 4C; Figure S6A)

### Validation of the THOR-1–SIF-1 interface *in vivo*

To validate the THOR-1–SIF-1 interaction interface defined by the crystal structure, we introduced targeted mutations into the endogenous *sif-1* locus using CRISPR/Cas9. We focused on three surface-exposed SIF-1 residues—Arg^142^, Thr^147^, and Arg^151^—identified in the structure as contributing to the binding interface. To disrupt these interactions, we introduced two charge-reversal mutations (Arg^142^Glu and Arg^151^Glu) and a Thr^147^Ala substitution, the latter removing the side-chain hydroxyl group and thereby abolishing its hydrogen bonding potential. Heterozygous individuals carrying these three mutations were viable and fertile; however, ∼25% of their progeny died during embryogenesis and only homozygous wt and heterozygous mutant individuals were viable and fertile (Figure S7). This phenotype is consistent with the complete loss of SIF-1 antidote activity.

To further dissect the contribution of individual SIF-1 residues to THOR-1 binding, we introduced substitutions at two of the key positively-charged residues: Arg^142^ and Arg^151.^ Charge-neutralizing mutations at either position—Arg^142^Ala or Arg^151^Ala—were largely tolerated, as homozygous lines carrying these mutations could be derived and propagated (Figure 4D). However, Arg^142^Ala and Arg^151^Ala mutants exhibited elevated developmental delay suggesting a partial (hypomorphic) loss of antidote function (Figure 4D). Consistent with this, the double charge-neutralizing mutant—Arg^142^Ala; Arg^151^Ala— was lethal, indicating that these residues act synergistically to support SIF-1 antidote activity (see methods; Figure S7). To determine whether this requirement reflects the presence of positive charge rather than specific side-chain identity, we substituted each arginine with lysine. Both Arg^142^Lys and Arg^151^Lys mutants were viable but showed a delayed phenotype similar to the charge-neutralizing mutations, suggesting that the specific chemical properties and geometry of the arginine side chains, rather than positive charge alone, contributes to productive THOR-1 recognition (Figure 4D). Arg^142^Asn and Arg^151^Gln substitutions were tolerated, although Arg^142^Asn produced the strongest phenotype among these viable variants, causing both increased lethality and developmental delay (Figure 4D). In contrast, charge reversal had substantially more severe consequences. Arg^142^Glu was lethal, as no homozygous animals could be recovered from heterozygous mothers, whereas Arg^151^Glu homozygotes were viable but exhibited high levels of embryonic and larval lethality (Figure 4D, Figure S7). Together, these results indicate that residues Arg^142^ and Arg^151^ make important, partially redundant contributions to SIF-1 antidote activity. The stronger phenotypes caused by charge-reversal mutations further support an electrostatic contribution to toxin recognition, while the residual activity of several charge-neutralizing substitutions indicates that the interface is not dependent on either arginine residue alone. Our genetic analysis thus provides *in vivo* validation of the THOR-1–SIF-1 interface identified structurally.

### Geographic distribution and recombination patterns of antagonistic TAs

The coexistence of antagonistic TAs raises important evolutionary questions: how are these mutually incompatible elements distributed in natural populations, and how do they shape chromosome evolution? Individual TAs are expected to benefit from tight linkage between toxin and antidote genes, as recombination can generate resistant antidote-only haplotypes. Conversely, simulations predict that recombination between antagonistic TAs can generate composite haplotypes carrying both elements, which may outcompete single-TA haplotypes^6^. Thus, the evolutionary dynamics of antagonistic TAs are likely to depend strongly on the local recombination landscape. To investigate the evolutionary dynamics in natural populations, we analysed sequencing data from 676 wild *C. tropicalis* isolates collected worldwide. We selected 622 reference isotypes from the *Caenorhabditis* Natural Diversity Resource (CaeNDR)^17^ and supplemented these with additional wild isolates sequenced in our laboratory (see Methods). We performed admixture analysis on LD-pruned variant calling data, with the number of hypothesized subpopulations K=9 chosen as optimal (see Methods). The resulting subpopulations broadly corresponded to the geographic distribution of the isolates. Consistent with Wang et al. (2026)^18^, we observed that Hawaiian and Taiwanese populations show higher within-group diversity, as evidenced by their clustering in multiple ancestry groups (Figure 5A). Next, for each isolate, their TA carrier status was inferred by aligning short-read sequencing reads to the two reference carrier genomes, NIC203 and EG6180 (see Methods).

**Figure 5.**
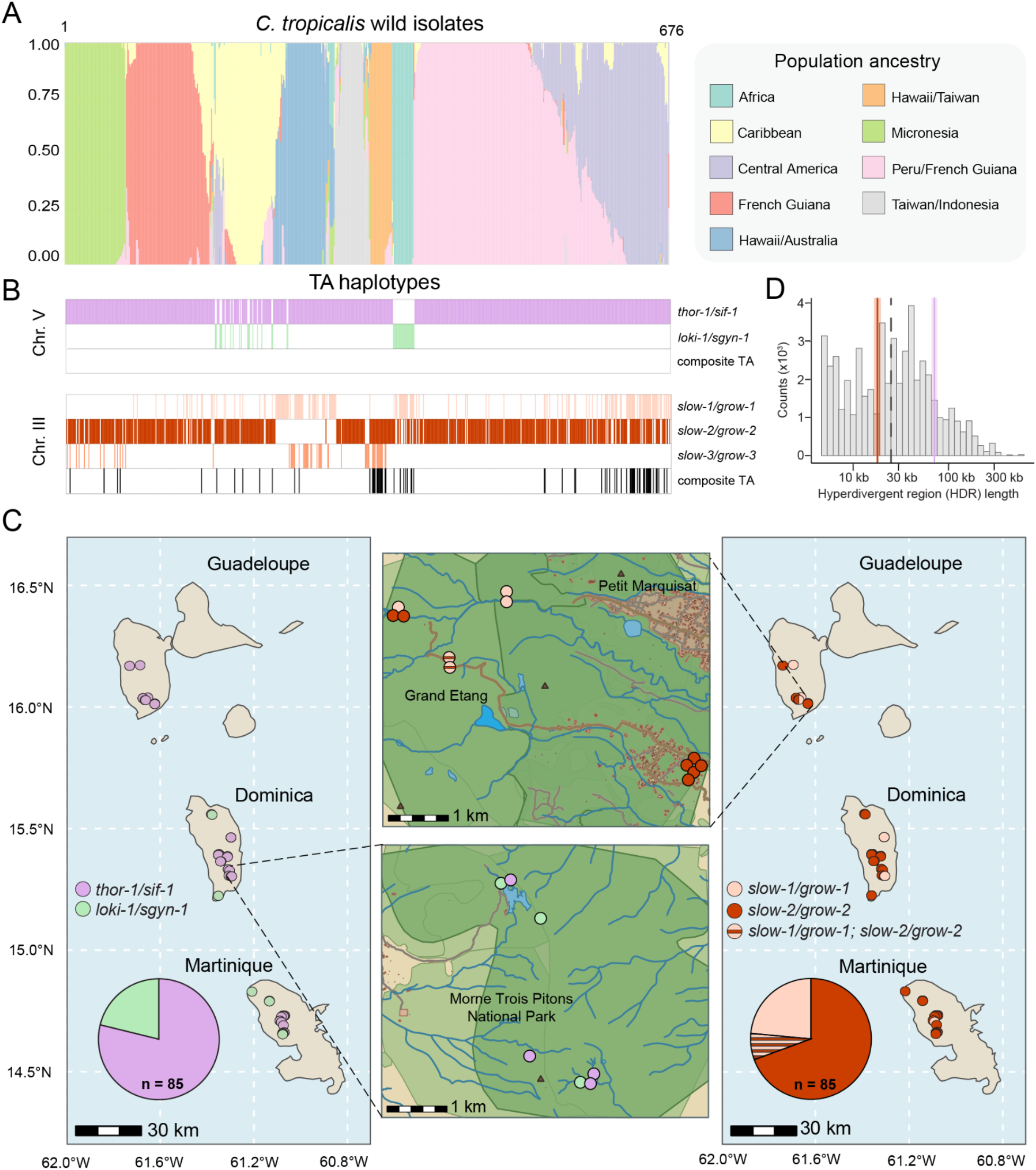
Geographic distribution of antagonistic TAs in *C. tropicalis* natural populations. **(A)** Population structure of 676 *C. tropicalis* wild isolates inferred using STRUCTURE. Each vertical bar represents one isolate, and the colored segments indicate its estimated membership proportions in each of the K inferred genetic clusters. Isolates are grouped according to geographic origin. Results are shown for K=9. **(B)** Antagonistic TA haplotypes carried by each of the 676 isolates, shown in the same order as in (B), for the antagonistic TAs on Chr. III and V. Composite haplotypes comprising two linked TA pairs are shown in black. For the *slow/grow* TA family, black lines indicate isolates carrying any combination of two linked *slow/grow* TA pairs. **(C)** Distribution of the two antagonistic TA haplotypes across Guadeloupe, Dominica, and Martinique. Pie charts indicate the relative frequency of each haplotype among 85 wild isolates sampled across the three Caribbean islands. **(D)** Size distribution of hyperdivergent regions (HDRs). The dotted line indicates the median HDR length. The purple vertical line denotes the HDR containing *thor-1/sif-1*, whereas the dark red vertical line denotes the HDR containing *slow-2/grow-2*.

We found that 93.5% (632/676) of all wild isolates carried the *thor-1/sif-1* TA, whereas the remaining 6.5% (44/676) carried the *loki-1/sgyn-1* TA (Figure 5B; Table S6). However, these frequencies are likely influenced by uneven geographic sampling. Most wild isolates in the collection originate from Central and South America and Southeast Asia (69%), regions where *thor-1/sif-1* predominates (96%) (Figure 5B). In contrast, isolates from Atlantic and Indian Ocean islands (including Capo Verde and La Réunion) almost exclusively carried *loki-1/sgyn-1* (94%), even though only 17 isolates from the region are available. Despite this uneven global distribution, the two TAs occur in close geographic proximity in nature. In both the Caribbean and Hawaiian islands, isolates carrying either *loki-1/sgyn-1* or *thor-1/sif-1* were collected from nearby locations (Figure 5C, Table S6, and Data S1). Most strikingly, we identified six pairs of isolates carrying opposing haplotypes that were collected at identical coordinates; five of these pairs were also collected on the same date (Table S7). These observations suggest that carriers of the antagonistic TAs encounter one another in natural populations and have opportunities to hybridize.

Yet, despite evidence of geographic overlap, we found no isolates carrying recombinant haplotypes that combined *thor-1/sif-1* with *loki-1/sgyn-1* (Figure 5B). One possible explanation is that the TAs occupy alternative haplotypes within a large hyperdivergent region (HDR). In selfing *Caenorhabditis* species, the relatively homogeneous genomic background is punctuated by highly divergent haplotype blocks that can differ extensively in nucleotide sequence, structural organization, copy number, and gene content^19,20^. We used the HDR annotation from CAENDR^17^ with NIC58 (*thor-1/sif-1* and *slow-2/grow-2* carrier) as a reference to investigate the potential influence of HDRs on antagonising TAs dynamics. The HDR containing *thor-1/sif-1* spans ∼ 71 kb, exceeding the length of 85% of all annotated regions and being 2.8-fold larger than a median HDR. If compared within carrier isolates, this block exceeds the length of ∼81% of HDRs and is 2.8-fold larger. We also annotated HDRs using EG6180 as a reference (see Methods). The median length of the HDR containing *loki-1/sgyn-1* is ∼119 kb, longer than 96% of HDRs and 10.8 times more than a median HDR. Within isolates, the numbers stayed consistent at 97% and 10.8 times, respectively. This indicates that *loki-1/sgyn-1* haplotype is not just present in isolates that are generally more distant from the reference and are expected to have more (and longer) HDRs, but also is more divergent compared to the rest of the carrier genome (Figure 5D). Although reduced recombination is not a defining feature of hyperdivergent regions, the extensive sequence and structural divergence between these alternative haplotypes may limit successful recombination in nature.

### A second family of antagonistic TAs forms composite TAs in nature

A second example of antagonistic TAs is represented by the *slow-1/grow-1* (NIC203) and *slow-2/grow-2* (EG6180) TAs in Chr. III. Unlike *loki-1/sgyn-1* and *thor-1/sif-1*, these two TAs are evolutionarily related: SLOW-1 and SLOW-2 are homologous toxins, and their respective antidotes, GROW-1 and GROW-2, are also homologous^8^. Despite this shared ancestry, their antidote activity remains highly specific, with each GROW neutralizing only its cognate SLOW toxin. Thus, the two homologous loci function as antagonistic TAs. Analogously to the analysis carried out for *loki-1/sgyn-1* and *thor-1/sif-1*, we surveyed the prevalence of both TA across the same panel of 676 wild *C. tropicalis* isolates (Figure 5B).

We could unambiguously assign either the *slow-1/grow-1* or *slow-2/grow-2* haplotype to 93% of the isolates; however, the remaining 46 isolates could not be assigned to either class, suggesting the presence of an additional haplotype. To investigate this apparent third haplotype, we focused on NIC1494, a wild isolate collected in Hsinchu County, Taiwan. We generated a high-quality genome assembly using long-read Nanopore sequencing followed by de novo assembly with Flye^21^. Analysis of the NIC1494 genome confirmed the existence of a third, distinct TA haplotype. The predicted toxin was more closely related to SLOW-1 than to SLOW-2 (64% vs 32% amino-acid identity), and was found in tight linkage with two antidote genes (Figure S8). One antidote shared 99% amino-acid identity with GROW-1, whereas the second exhibited 93% identity to GROW-1 and represented a previously undescribed antidote variant (Figure S8). We therefore designated this haplotype *slow-3/grow-1;grow-3*, hereafter referred to as *slow-3/grow-3* for simplicity.

Analysis of the three existing haplotypes revealed a markedly different evolutionary pattern to *loki-1/sgyn-1* and *thor-1/sif-1* (Figure 5B and Figure 6A). Notably, short-read alignments predicted that 10.4% of isolates (70/676) simultaneously carried two of the three *slow* haplotypes, with *slow-2* co-occurring with either *slow-1* or *slow-3* (Figure 5B and Table S6). Because the individual *slow/grow* haplotypes occupy the same locus on Chr. III, this pattern could be explained either by recombination generating composite TA haplotypes containing two linked TAs, or by translocation of one slow/grow locus to a different genomic location, allowing the two systems to segregate independently. To distinguish between these possibilities, we examined publicly available *de novo* genome assemblies from strains predicted to carry two TA haplotypes. This analysis strongly supported the first scenario. For example, the reference strain JU1373, isolated from La Réunion Island, carries linked copies of both *slow-1/grow-1* and *slow-2/grow-2* on Chr. III only ∼15kb part (Figure 6B). Analysis of the recombinant haplotype sequence revealed that *slow-2* is likely inactive in JU1373 due to the presence of a deleterious frameshift mutation but all other genes are still functional (Figure 6C). To test whether both antidotes remain functional and whether *slow-1* retains toxin activity in the composite JU1373 haplotype, we performed reciprocal crosses between the JU1373 and EG6180 strains. When JU1373 hermaphrodites were mated to EG6180 males, we observed 21% developmental delay and larval arrest among their F_2_ progeny (n=200) (Figure 6C). Consistent with an active JU1373 *slow-1/grow-1* element, F_2_ EG/EG homozygotes were largely affected (87.8% delay; n=41), as these individuals lacked the corresponding *grow-1* antidote. By contrast, only 2.4% of JU/JU homozygotes were delayed (n=42), indicating that the JU1373 *grow-2* antidote remains functional and protects against the EG6180 SLOW-2 toxin (Figure 6C).

**Figure 6.**
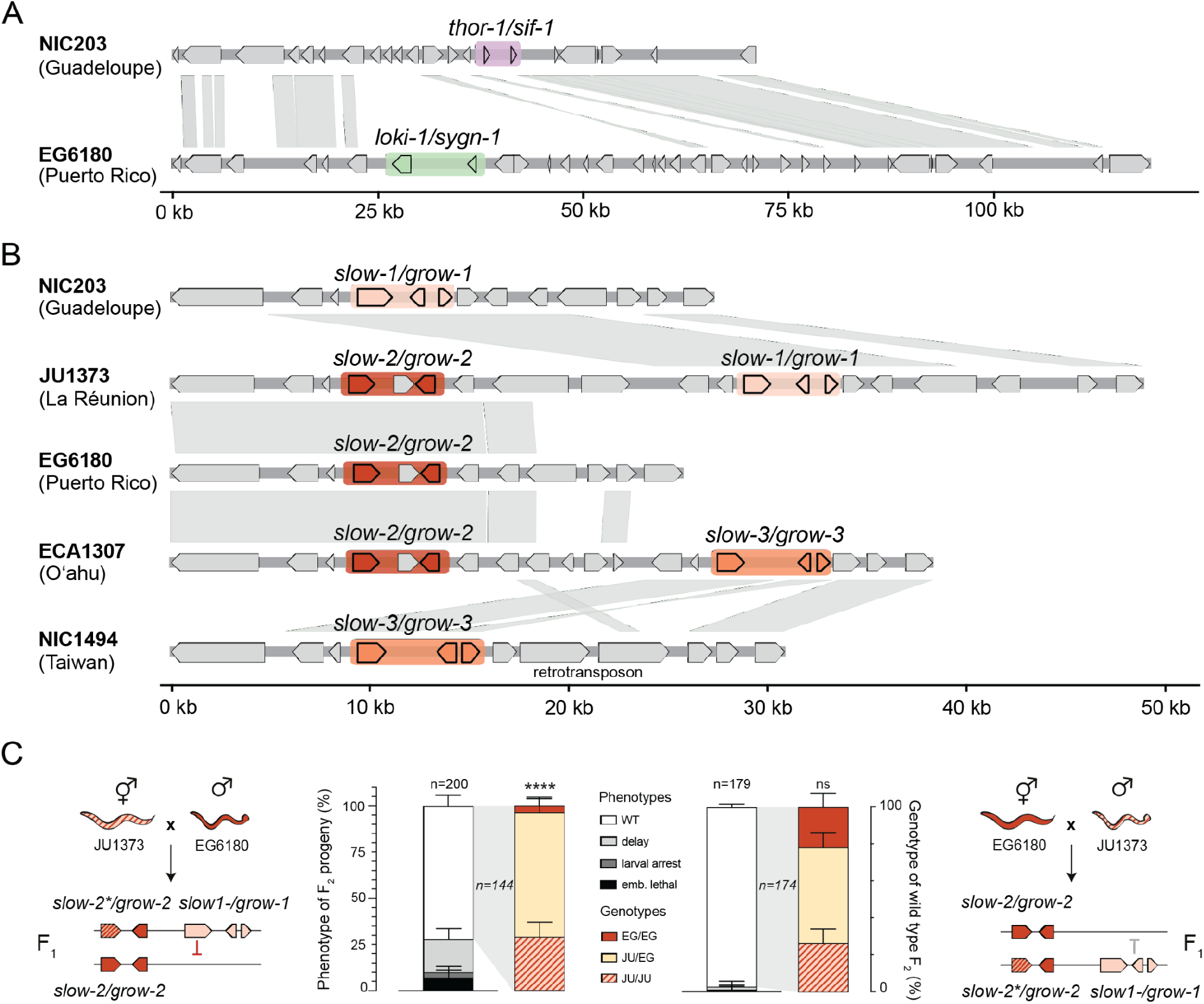
Antagonistic TA loci exhibit contrasting patterns of recombination in natural populations. **A)** Alignment of the two alternative TA haplotypes and their flanking regions in Chr. V: NIC203, carrying *thor-1/sif-1*, and EG6180, carrying *loki-1/sgyn-1*. **(B)** Alignment of the *slow/grow* TA haplotypes and their flanking regions on Chr. III. The three individual TA haplotypes are represented by NIC203 (*slow-1/grow-1*), EG6180 (*slow-2/grow-2*), and NIC1494 (*slow-3/grow-3*). Also shown are two composite haplotypes containing two linked TAs: JU1373 (*slow-1/grow-1* and *slow-2/grow-2*) and ECA1307 (*slow-2/grow-2* and *slow-3/grow-3*). In both panels, gray shading indicates regions with >80% nucleotide sequence identity. **(C)** Reciprocal crosses between JU1373 and EG6180 used to test the activity of the composite *slow-1/grow-1* and *slow-2/grow-2* TA haplotype in JU1373. JU1373 hermaphrodites were crossed with EG6180 males (left), and EG6180 hermaphrodites with JU1373 males (right). The JU1373 *slow-2* toxin is predicted to be inactive due to a deleterious mutation. *P* values calculated using exact multinomial goodness-of-fit test to the expected Mendelian ratio.

In the reciprocal cross, both JU/JU and EG/EG F2 homozygotes were predominantly wild type, and the wild-type F_2_ genotype distribution was not different from Mendelian expectation (*P = 0.66*; n = 174; Figure 6C). The absence of toxicity toward EG/EG progeny when the JU1373 haplotype was inherited paternally indicates that JU1373 *slow-1/grow-1* is subject to epigenetic regulation, consistent with the epigenetic licensing previously described for the NIC203 *slow-1/grow-1* element.^8^ Importantly, the *slow-2* toxin is not inactivated in all recombinant haplotypes carrying both *slow-1/grow-1* and *slow-2/grow-2*. We identified additional composite haplotypes in which both toxin genes are predicted to remain functional, including those carried by the Indonesian isolates HPT33 and HPT56.

Analysis of the long read sequencing data of the wild isolate ECA1307, collected on Oʻahu, Hawaiʻi, provided independent evidence for a second composite TA haplotype, comprising linked *slow-2/grow-2* and *slow-3/grow-3* TAs (Figure 6B). Both *slow-2* and *slow-3* are predicted to encode functional toxins, and all associated antidotes appear intact. Notably, this haplotype is predicted to be resistant to all previously identified *slow/grow* haplotypes, as it encodes the full repertoire of antidotes—*grow-1*, *grow-2,* and *grow-3*. Although we found no evidence for a strain carrying all three *slow/grow* TAs, short-read alignments also identified isolates consistent with a composite *slow-1/grow-1*; *slow-3/grow-3* haplotype. However, the absence of a corresponding genome assembly prevented us from confirming this putative recombinant haplotype. Assuming that all isolates lacking genome assemblies carry either single or composite TA haplotypes analogous to those validated above, we estimate that 51 isolates (7,5%) harbor a composite *slow-1/grow-1; slow-2/grow-2* haplotype, 17 (2.5%) harbor a composite *slow-2/grow-2; slow-3/grow-3* haplotype, and 2 (0.3%) harbor a putative composite *slow-1/grow-1; slow-3/grow-3* haplotype.

Unlike *thor-1/sif-1* and *loki-1/sgyn-1*, the hyperdivergent region containing the *slow* locus is comparatively short, with a median length of ∼18 kb. This is longer than only 36% of annotated hyperdivergent regions and corresponds to 0.7 times the median hyperdivergent block length (Figure 5D). Furthermore, sequence divergence is largely restricted to the fast evolving TA genes and their immediate genomic neighbourhood, while flanking regions remain conserved (Figures 6A and 6B). These observations reveal two contrasting evolutionary trajectories for antagonistic TAs. *thor-1* and *loki-1* originated independently and occupy distinct hyperdivergent haplotypes with a largely non-homologous genomic background. In contrast, *slow-1*, *slow-2*, and *slow-3* represent divergent alleles of the same ancestral TA that share conserved genomic context and flanking regions. This common genomic background allows recombination and formation of haplotypes carrying multiple *slow* alleles. Together, these findings suggest that the evolutionary fate of antagonistic TAs is strongly influenced by their origin and genomic context: independently evolved elements remain locked in conflict, while allelic variants can generate composite TA haplotypes capable of driving in a population and more complex population dynamics.

## Discussion

We genetically dissected two selfish TAs in *C. tropicalis* which are in direct molecular conflict: *thor-1/sif-1* and *loki-1/sgyn-1*. These elements occupy alternative haplotypes within hyperdivergent regions and act against one another in heterozygous crosses. Our findings explain earlier observations by Noble et al. who studied crosses between a different pair of wild isolates^7^. In a recombinant inbred line panel derived from JU1373 (La Réunion) and NIC58 (French Guiana), they detected strong transmission distortion and persistent heterozygosity on the right arm of chromosome V. Follow-up crosses suggested the presence of antagonistic TAs, but the underlying genes remained unknown^7^. NIC58 carries *thor-1/sif-1*, whereas JU1373 carries *loki-1/sgyn-1* (Table S6), suggesting that antagonism between these elements could account for both phenomena.

Our results also provide an alternative explanation for the differences in TA activity observed between reciprocal crosses of NIC58 and JU1373. Noble and colleagues reported that the NIC58 TA was active in the presence of NIC58 mitochondria but largely inactive in animals carrying JU1373 mitochondria, whereas the JU1373 TA was active in reciprocal crosses^7^. In contrast, our findings indicate that the asymmetry can be explained by epigenetic licensing of *thor-1*. When *thor-1/sif-1* is maternally inherited, maternally deposited *thor-1* transcripts license the zygotic allele and protect it from piRNA-mediated repression, as previously shown for *slow-1*^8^. When the TA is inherited paternally, the absence of maternal *thor-1* transcripts results in germline silencing of *thor-1* in F_1_ hermaphrodites and, consequently, reduced toxicity in their F_2_ progeny. Thus, the parent-of-origin effects previously attributed to mitochondrial-nuclear interactions can instead be explained by epigenetic licensing via maternal transcripts.

Recent studies have identified TA elements underlying genetic incompatibilities in diverse taxa, including fungi, nematodes, insects, plants, and mice^2^. While only a handful of TAs have been characterized in most species, the seven now identified in *C. tropicalis* provide an unusually rich system for uncovering general principles of TA evolution. One emerging theme is the recurrent recruitment of F-box proteins as antidotes. Four *C. tropicalis* antidotes—SGYN-1, KSS-1, KSS-2, and KSS-3^9^—are F-box proteins, as are PHA-1 in *C. elegans*^9^ and SHLS-2 in *C. nigoni*^22^, indicating that this protein class has been independently co-opted as an antidote in *C. tropicalis* and in two other nematode species. Our phylogenetic reconstruction further suggests that SIF-1 also evolved from an F-box protein but subsequently lost its F-box domain. The crystal structure of the THOR-1–SIF-1 complex provides direct molecular insight into the functional consequence of this evolutionary change. Rather than functioning as an SCF ubiquitin ligase through an F-box domain to promote toxin degradation, SIF-1 directly engages THOR-1 through an extensive protein–protein interface. These findings suggest that loss of the F-box domain was accompanied by a shift from ubiquitin-dependent toxin degradation to direct toxin neutralization.

The remaining SIF-1 structure appears to provide a dedicated surface for THOR-1 recognition, consistent with an evolutionary repurposing of an ancestral F-box protein scaffold. Notably, the THOR-1-interacting surface of SIF-1, including the structural element comprising residues 100–102, is poorly conserved among SIF-1-related proteins, whereas the opposite surface is highly conserved (Figure 6B). This contrasting conservation pattern is consistent with diversification of the toxin-binding surface while other structural features of SIF-1 have remained constrained. The conserved opposite surface may therefore have been retained for a distinct, as-yet-uncharacterized function. The structure does not, however, establish the precise molecular consequence of THOR-1 binding. Because the cellular target of THOR-1 remains unknown, it is not yet possible to determine whether SIF-1 neutralizes the toxin by directly occluding its target-binding surface, by inducing an inhibitory conformational state, or through another mechanism.

Although we now have a clearer molecular understanding of antagonistic TAs, their consequences for population dynamics and long-term evolution remain poorly understood. While laboratory crosses with antagonistic TAs generate an excess of heterozygotes, population-genetic simulations indicate that this does not necessarily promote the stable coexistence of alternative haplotypes (balancing selection)^23^. Instead, particularly under high selfing, antagonistic TAs can act as very effective barriers to gene flow: once one TA is established, even a weakly penetrant toxin can prevent the invasion of an alternative TA with substantially higher penetrance^23^. This prediction is consistent with our observation that alternative antagonistic TA haplotypes can occur in close geographic proximity in nature, while remaining locally segregated. To examine this at a much finer spatial scale, we analyzed an independent set of 39 isolates collected along a 12-m forest path in French Guiana, but found that all carried the same TA haplotype (Table S6). Our large-scale survey therefore provides a framework for targeted sampling in regions of haplotype overlap to determine the spatial scale over which antagonistic TAs may restrict gene flow.

Existing models, however, do not capture several features revealed here, including epigenetic licensing and the formation of composite TA haplotypes^23^. Because a paternally introduced, unlicensed toxin can remain epigenetically repressed for several generations, the success of an invasive TA may depend not only on its penetrance and frequency but also on its direction and history of transmission. Composite *slow/grow* haplotypes introduce a further asymmetry. Their evolution is unlikely to reflect increased drive strength, because tight linkage largely eliminates any advantage from carrying already highly penetrant toxins. Instead, composite haplotypes may be favored because their expanded antidote repertoire protects them against alternative TAs, thereby increasing resistance to invasion. The naturally occurring composite haplotypes identified here therefore illustrate how recombination can transform competing TA systems into multi-antidote haplotypes with the potential to resist multiple toxin variants.

Thus, in addition to their selfish drive, antagonistic TAs may shape population structure through three interacting processes: i) the formation of localized barriers to gene flow, ii) selection for piRNAs mediating recognition and silencing of toxins, and iii) selection for antidote-mediated immunity to competing selfish elements via recombination. At the molecular level, our structural analysis further shows that this conflict can drive the evolution of new toxin-recognition mechanisms: an ancestral F-box antidote can lose its canonical degradation machinery and become a direct protein–protein interaction module specialized for toxin neutralization. Together, these findings illustrate how antagonistic selfish elements can generate evolutionary innovation at the levels of protein structure, gene regulation, and chromosome organization.

## Acknowledgments

Research in the Burga lab is supported by the Austrian Academy of Sciences, the city of Vienna, and a European Research Council (ERC) Starting Grant under the European Union’s Horizon 2020 program (ERC-2019-StG-851470), and the Stand-Alone Austrian Science Fund (FWF) grant P34880 (G.D. and A.B.). A.Koreshova is supported by a Boehringer Ingelheim Fonds (BIF) PhD Fellowship. C.B. is supported by the CNRS (Centre National de la Recherche Scientifique). Library preparation and sequencing were performed at the Vienna BioCenter Core Facilities (VBCF) NGS Facility. *C. tropicalis* transgenic lines were generated by the IMBA Fly and Worm Injection Facility. Insect cell expression and SEC-MALS experiments were performed by the Protein Technologies Facility at VBCF. C.B. thanks the CNRS Nouragues Research Station in French Guiana for hosting and supporting the fieldwork.

## Data availability

All raw sequencing data and genome assemblies are available under National Center for Biotechnology Information (NCBI) project no. PRJNA1524375. The crystal structure of the THOR-1–SIF-1 complex has been deposited in the Protein Data Bank (PDB) under accession code 35AW.

## Author contributions

A.B. and A.Koreshova conceived and developed the project. A.Koreshova and P.T. performed genetic crosses, TA fine-mapping, and molecular genetics experiments. A.Koreshova performed genomic analyses of natural isolates. C.B. did field-work and isolated natural *C. tropicalis* strains. T.L. performed phylogenetic analyses. A.H. purified and characterized the THOR-1–SIF-1 complex. A.Kara and G.K. performed crystallography and structure determination under the supervision of G.D. G.D. carried out structural analyses, PDB structure deposition, and prepared figures. A.Koreshova and A.H. performed in vivo validation of structure-guided predictions. J.G. and P.D. designed and generated transgenic worm lines. A.B., G.D., and A.Koreshova wrote the manuscript.

## Material and Methods

### Maintenance of worm strains

Nematodes were maintained on modified NGM plates supplemented with 1% agar and 0.7% agarose, a formulation used to minimize burrowing behavior in *C. tropicalis*. All cultures were fed *Escherichia coli* OP50. Unless otherwise noted, *C. tropicalis* strains were propagated at 25°C. To eliminate bacterial contamination, gravid animals were subjected to standard alkaline hypochlorite treatment. Briefly, 500 µL of a freshly prepared 1:1 mixture of 1 M NaOH and 5% sodium hypochlorite was added to 750 µL of worm suspension in M9 buffer. Following dissolution of adult carcasses, embryos were recovered and the bleaching reaction was terminated by two successive washes in 14 mL M9 buffer. Some strains were provided by the CGC, which is funded by NIH Office of Research Infrastructure Programs (P40 OD010440). A complete list of strains used in this study is provided in Table S8.

### Phenotyping and genotyping of crosses

Crosses were established by placing 4–5 L4-stage hermaphrodites together with 25–50 males (approximately 1:6–1:10 hermaphrodite-to-male ratio) on 5-cm plates containing modified NGM medium. After 48 h, fifteen F_1_ L4 progeny were transferred as a group to fresh plates and isolated individually the following morning to synchronize production of the F_2_ generation. Following 2–3 h of egg laying, F_2_ hermaphrodites were collected for PCR-based genotyping. On the same day, at least ten embryos from each genotyped heterozygous F_1_ animal were transferred to separate plates for phenotypic analysis. F_2_ progeny were monitored daily until death or attainment of reproductive maturity. Phenotypes scored included embryonic and larval lethality, developmental arrest, delayed reproductive development, and sterility. For genotyping, individual worms were lysed in a buffer containing 10 mM Tris-Cl pH8, 50 mM KCl, 2.5 mM MgCl_2_, 200 µg/ml Proteinase K. PCR amplification was carried out using an in-house hot-start Taq polymerase preparation (Molecular Biology Service, IMBA). When required, PCR amplicons were verified by Sanger sequencing. Genotyping success was reduced for animals that arrested during embryogenesis or early larval development. For the cross of JU1373 males to EG6180 hermaphrodites, dumpy worms were used instead of WT EG6180 to simplify the screening for heterozygous F1 progeny due to low mating efficiency of JU1373 males (Figure 6C). Additional details regarding crossing schemes and raw data for all genetic crosses are provided in Table S9.

### Generation of transgenic lines

Genome editing was performed using CRISPR–Cas methodologies adapted from previously published protocols. Cas9 or Cas12a nuclease (IDT; 250 ng/µL) was preassembled with guide RNAs consisting of 200 ng/µL crRNA and 333 ng/µL tracrRNA (IDT) by incubation at 37°C for 10 min. Subsequently, a fluorescent co-injection marker plasmid (2.5 ng/µL; pCFJ90-mScarlet-I or pCFJ90-mNG for edits in mScarlet-tagged backgrounds) was added to the injection mixture. For homology-directed repair, either donor oligonucleotides (200 ng/µL) or biotinylated, denatured PCR-derived donor templates (100 ng/µL) were included. Injection mixes were introduced into young adult hermaphrodites, and fluorescent F1 progeny were isolated individually. Successful genome edits were identified by PCR screening and confirmed by Sanger sequencing of candidate lines. To generate single-copy transgene insertions, we adapted a split-selection strategy originally developed for *C. elegans*. A synthetic landing pad (SLP) carrying a split hygromycin-resistance cassette was first integrated on chromosome IV of the *C. tropicalis* strain EG6180. Repair templates consisted of plasmids encoding the desired insert together with the N-terminal fragment of the hygromycin-resistance gene driven by the ribosomal promoter Ctr-rps-20p and were injected at 50–60 ng/µL. Hygromycin B selection was initiated 3–4 days after injection by adding 600 µL of a 5 mg/mL solution to each plate. Resistant animals surviving 7–10 days after selection were isolated, propagated, and screened by PCR and Sanger sequencing to verify correct transgene integration. Sequences of all guide RNAs, HDR donor templates, and primers used for SLP-mediated integrations are provided in Table S10.

### Fine mapping of the *loki-1/sgyn-1* and *thor-1/sif-1* TAs

To narrow down the introgression region, the crosses of 4 Chr. V NIL hermaphrodites to 40 NIC203 males were performed, and F2 progeny from heterozygous F1 mothers was singled. PCR by three pairs of primers covering indel loci spread along the introgression region was used to screen for recombination events (Table S9). Homozygous lines were established from individual worms that had an indel pattern different from both NIC203 and Chr. V NIL, and sequenced with Illumina MiSeq to establish the precise coordinates of the new introgression regions. The resulting sub-NILs were crossed to Chr. V NIL males to examine whether they carry the TA from EG6180 (with expectation of no larval arrest in F2 since both lines carry an antidote), NIC203 (with expectation of embryonic lethality in F2), or both. The regions that must contain both TAs were deduced from the resulting genomic and phenotypic patterns. The genes in the regions of interest were analysed using BLAST and RNA-seq data to search for pairs of genes that are 1) divergent or absent compared to the non-carrier strain, 2) have detectable expression, and 3) are in close proximity. Verification of the candidates was performed using Crispr/Cas genomic editing to create a null allele.

### Expression and purification of the THOR-1–SIF-1 complex

N-terminally 6x-His-tagged SIF-1 and untagged THOR-1 were expressed from a baculovirus expression vector pGB-Dest in Sf9 cells at 27 °C. The cells were harvested 3 days post infection and stored at −70 °C until purification. Cell pellets corresponding to 0.5 L of medium were thawed on ice, resuspended in 40 mL lysis buffer (2x PBS, 20 mM imidazole, 0.04 µg/ml Benzonase (final concentration; from a 0.4 mg/ml stock (Molecular Biology Service, IMBA)), EDTA-free cOmplete protease inhibitor cocktail (Roche), 1 mM TCEP, 0.01% Triton X-100), incubated on a tube roller for 30 minutes at 4 °C and centrifuged at 50,000 rcf at 8 °C. Affinity purification was performed on a HisTrap HP 5 mL (Cytiva) with 2 wash steps of 20 CV 40 mM imidazole, then 10 CV 60 mM imidazole and proteins were eluted by a linear gradient over 20 CV to 500 mM imidazole. Fractions containing both proteins of interest were pooled and diluted 1/10 into 20 mM Tris pH7.5, 10 mM NaCl, 0.2 mM TCEP for ion exchange chromatography on ResourceQ 6 mL (Cytiva). The protein complex was eluted by a linear gradient of NaCl and fractions containing the pure complex were pooled and concentrated. The concentrated complex was then applied to a size exclusion chromatography step on a Superdex200 10/300 Increase (Cytiva) in 20mM Tris pH7.5,150mM NaCl, 0.2mM TCEP. The concentrated fractions after this step and the same buffer were used for the SEC-MALS experiment, for which the sample was diluted to 1 mg/ml and was run in triplicate.

### In situ proteolysis and crystallization

The purified THOR-1–SIF-1 complex was subjected to limited proteolysis with trypsin, chymotrypsin, GluV8, and thermolysin to identify stable proteolytic fragments. Thermolysin was finally chosen for in situ proteolysis crystallization based on the well-defined and relatively stable digestion products observed in the proteolysis screen. For crystallization, the purified THOR-1–SIF-1 complex (12 mg/ml) was mixed with thermolysin at a final concentration of 1.2 µg/ml and directly subjected to crystallization trials using a variety of commercial crystallization screening kits at either 22°C or 4°C. Single crystals suitable for X-ray diffraction were obtained after 3–5 days at 4°C in a condition containing 0.1 M sodium acetate, 0.1 M Tris-HCl (pH 8.5), and 30% (w/v) PEG 8000.

### X-ray data collection and structure determination

X-ray diffraction data were collected at 100 K on beamline ID30B at the European Synchrotron Radiation Facility (ESRF) using a wavelength of 2.0664 Å. A total of 1,800 diffraction images were collected with an oscillation range of 0.35° per image. The diffraction data were indexed, integrated, and scaled using XDS^24^. The crystals belonged to space group P2₁2₁2, with unit-cell parameters a = 90.13 Å, b = 98.48 Å, and c = 40.46 Å. The dataset extended to 2.20 Å resolution, with an overall completeness of 99.36% and a multiplicity of 11.99.

The structure of the THOR-1–SIF-1 complex was determined by single-wavelength anomalous diffraction (SAD), exploiting the anomalous signal from sulfur atoms, using AutoSol^25^, as implemented in the PHENIX suite^26^. The automatically built model was subsequently inspected and manually adjusted using Coot^27^, followed by iterative refinement using phenix.refine^28^. The final model was refined against data extending from 20 to 2.20 Å resolution and contained 2,357 protein non-hydrogen atoms and 192 solvent atoms. The final R-work and R-free values were 0.187 and 0.231, respectively. Model geometry was assessed using standard crystallographic validation procedures; 95.7% of residues were in favored Ramachandran regions, 4.3% were in allowed regions, and no residues were classified as outliers (Table S5). The final coordinates have been deposited in the Protein Data Bank (PDB) under accession code 35AW.

### Identification of SIF-1 and SGYN-1homologs and phylogenetic reconstruction

SIF-1 and SGYN-1 protein structures were predicted using AlphaFold3^29^, and the best-ranked model was used as the reference structure. Protein structure databases were generated from 50 nematode genomes using Foldseek v10^30^ createdb with default settings. We searched the reference structure against these databases using the Foldseek easy-search workflow with default settings. Candidate proteins were retained with an E-value <1e-2 and both target and query coverage ≥40%. The structure of the candidate proteins were predicted using the AlphaFold2 implementation in ColabFold v1.5.2^31^. Three models were generated using --num-recycle 3 --num-models 3, and the best-ranked model was selected for subsequent analyses. Each candidate structure was aligned to the reference structure using USalign v20241108^32^. Candidates with more than 30% of aligned residues had a distance <5 Å were retained for subsequent analysis. Pairwise structural similarities among candidate proteins were calculated using USalign^32^ in fast mode (-fast). The mean TM-score from each pairwise alignment was used to construct a pairwise protein distance matrix. We used NetworkX v3.1^33^ to construct a fully connected network from the distance matrix. Series of increasing distance cutoffs were applied to identify connected components, and the cutoff corresponding to the largest change in network connectivity was selected as the optimal threshold. The component containing the reference protein was retained, resulting in a final set of homologs. The protein sequences of the homologs were aligned using FoldMason v4^34^ with the easy-msa workflow and --refine-iters 100. The resulting multiple sequence alignment was used for phylogenetic reconstruction using IQ-TREE v2.1.3^35^.

### De novo protein structure and interaction prediction

Protein structure predictions were generated using AlphaFold3^29^. Protein–protein interaction predictions were also performed with AlphaFold 3 for SGYN-1 in combination with each annotated *C. tropicalis* SKR protein (Table S4).

### Phenotyping SIF-1 mutants

For the lines that were successfully established as homozygotes, phenotyping was performed as follows: wild-type looking L4 hermaphrodites were picked from a not starved plate, let develop into adults, transferred to a fresh plate and let lay for an hour. The eggs from this plate were singled and monitored daily until death or the plate being starved. For the lines that were not homozygous viable, progeny of heterozygous mothers was picked and monitored for embryonic lethality the next day. The survivors were propagated and genotyped. The procedure was repeated with subsequent heterozygous generations to increase the sample size.

### Short read sequencing, variant calling and admixture analysis

Total DNA was extracted from 1 freshly starved plate with Masterpure Complete DNA&RNA extraction kit according to the manufacturer’s protocol, and libraries were prepared using an in-house Tn5-based protocol and sequenced with NovaSeq SP (Illumina) in paired-end read mode by Next Generation Sequencing Facility at Vienna BioCenter Core Facilities (VBCF), member of the Vienna BioCenter (VBC), Austria. For a broad survey of natural variation in *C. tropicalis*, we analyzed 54 wild isolates from the Braendle collection that were not represented in CeNDR, together with publicly available sequencing data from 622 reference isotypes downloaded from CeNDR^17^. For the fine-scale survey, we sequenced 39 wild isolates collected in October 2021 at the Pararé research camp in the Nouragues National Nature Reserve, French Guiana. Nematodes were recovered from individual native fig substrates using Baermann funnel extraction immediately after collection^36^. along a single man-made path through the forest, covering an area approximately 12 m long and 0.5 m wide, with individual figs spaced approximately 30 cm apart. This sampling design was intended to assess whether distinct genotypes could colonize the same fruit substrate. When multiple nematode individuals were recovered from a single fig, an independent line was established from each individual. Illumina short reads were aligned to both NIC203 and EG6180 genomes with bwa mem v0.7.17^37^, and the variants were called with bcftools v1.9^38^. The vcf was filtered to include samples that have > 80% of sites genotyped, and only biallelic sites with > 90% genotyped and r2 < 0.5. The resulting vcf was converted to bed using plink v1.9^39^ and admixture v1.3^40^ was run 10 times for K values from 5 to 15. K=9 was chosen because it yielded the minimum cross-validation (CV) error among the tested values in NIC203.

### Annotation of Hyperdivergent regions and identification of TA haplotypes

The hyperdivergent regions annotation was performed largely as in Wang et al. (2026)^41^. The snp density was calculated from the vcf files described above with vcftools v0.1.16^42^. Percentage of bases covered at least at coverage = 1 was calculated using mosdepth v0.2.9^43^. Both procedures used a 1 kb window. The bins with variant count of 9 or larger or percent bases covered under 90% were classified as hyper-divergent, genome wide, across all isolates. Proximal HDRs separated by less than 5 kb were merged.

For identification of TA haplotypes, mean whole genome coverage and coverage at known toxin loci was calculated for each isolate aligned to both NIC203 and EG6180 with mosdepth v0.2.9^43^. Based on toxin coverage / total genome coverage percentage histograms, an initial cut-off of 50% was chosen as it was separating the two modes of the distribution representing carriers and non-carriers. It was subsequently used for *thor-1/sif-1* and *loki-1/sgyn-1* assignment. For *slow* allele assignment at this threshold, 46 isolates showed no *slow* allele present. The isolate with lowest coverage percentage on both *slow-1* and *slow-2*, NIC1494, was sequenced with Nanopore, and its genome was assembled as described in the section below. BLASTN search revealed a new *slow-3* allele in NIC1494. To identify *slow* alleles carried by different isolates and overcome their partial sequence similarity, all isolates were aligned to a panel containing all *slow* alleles and a set of conserved genes of NIC203 for the baseline coverage estimate. The conserved genes were selected from busco nematoda odb12 set^44^. BLASTP search against NIC203 and EG6180 proteins with -qcov_hsp_perc 80 -evalue 1e-10 thresholds was performed for all proteins from the busco database ranging from 300 to 500 aa in length. For the ones that had a single significant hit in both NIC203 and EG6180, reciprocal BLASTP search was performed to verify that these *C. tropicalis* proteins are one-to-one homologues of busco reference proteins. Only pairs of busco/NIC203 proteins that were in range of 90% to 110% of length of each other were kept to ensure there were no incomplete protein annotations in NIC203. 30 longest genes (all above 1500 bp) were selected for a trial alignment, and five were subsequently removed due to coverage more than 2x of the median coverage across the set, possibly due to presence of repetitive sequences. A fasta file containing the remaining 25 conserved genes together with *slow* alleles was used as a reference for the alignment of all isolates. A cut-off of 15% was chosen in the similar way as before and the borderline cases were examined visually in IGV^45^ to ensure continuous alignment of all detected *slow* alleles. The data processing was done in R^46^, all scripts are available upon request. The interactive map of isolates and their corresponding TAs was created with help of sf^47^ and leaflet^48^ packages and is available as Supplementary Data 1.

### Long read sequencing and genome assembly

Total DNA was extracted using the phenol-chloroform method. Libraries were prepared using no-shearing ligation protocol and sequenced on PromethION (Nanopore) by Next Generation Sequencing Facility at Vienna BioCenter Core Facilities (VBCF), member of the Vienna BioCenter (VBC), Austria. The reads were trimmed with cutadapt v1.18^49^ and assembled using flye v2.7.1^50^ with –nano-raw setting and estimated genome size of 80 Mb. The resulting assembly of NIC1494 is 81 955 248 bp long, N50 = 6, 98.4% busco^51^ complete. For ECA1307, unassembled Hi-Fi sequencing data was downloaded from SRA (run SRR36938953) and aligned to *slow-2* and *slow-3* genomic sequences with minimap2^52^. Successfully aligned reads were assembled using flye^50^ with --pacbio-raw setting, resulting in a single contig of 44 491 bp.

## Supplemental Information

Supplemental Figures S1-S8

Supplemental Tables S1-S10

Data S1

**Figure S1. Phenotypic characterization of null mutant lines.** Neither single-toxin mutants nor double toxin–antidote mutant combinations exhibited developmental defects above background levels. These results indicate that the corresponding genes are dispensable for normal *C. tropicalis* embryonic development, consistent with their proposed roles as selfish genetic elements.

**Figure S2. The *loki-1* toxin is homologous to *C. elegans ucr-2.3* but is highly divergent. (A)** Protein sequence alignment of *C. tropicalis* LOKI-1, the *C. tropicalis ucr-2.3* ortholog ORF019193, and *C. elegans* UCR-2.3. **(B)** AlphaFold3 prediction of the LOKI-1 protein structure. The overall confidence of the predicted model is very low, providing limited insight into the potential mechanism of LOKI-1 toxicity.

**Figure S3. Phylogenetic relationships among SGYN-1, SIF-1, and the SSLP family of F-box proteins and their genomic distribution. (A)** Phylogenetic tree including all annotated SSLP (SIF- and SGYN-like protein) family of intact F-box proteins and proteins that lost their F-box domains like SIF-1 from the *C. tropicalis* wild isolates NIC203 and EG6180 and from *C. brenneri*, together with the antidotes SGYN-1 and SIF-1. **(B)** Genomic distribution of SSLP genes in NIC203 (right) and EG6180 (left). In both isolates, SSLP genes form clusters on the right arm of chromosome V.

**Figure S4. Flexible regions in THOR-1 and SIF-1 and limited proteolysis of the THOR-1–SIF-1 complex. (A)** Predicted intrinsically disordered regions in THOR-1 (top) and SIF-1 (bottom), generated using IUPred2. **(B)** Limited proteolysis of the THOR-1– SIF-1 complex with increasing concentrations of thermolysin. Reproducible stable proteolytic fragments were observed. Proteins were visualized by Coomassie staining.

**Figure S5**. **Crystallization and X-ray diffraction of the THOR-1–SIF-1 complex**. **(A)** Representative crystal of the THOR-1–SIF-1 complex (left). Crystallographic data collection statistics (right). Space group, unit-cell parameters, completeness, resolution range, and Rmerge are shown for the THOR-1–SIF-1 crystal dataset. Inner- and outer-shell statistics correspond to 97.7–9.0 Å and 2.25–2.18 Å, respectively. **(B)** Representative X-ray diffraction image showing Bragg reflections extending to approximately 2.2 Å resolution.

**Figure S6. Structural details of the THOR-1–SIF-1 interaction interface. (A)** Residues in THOR-1 and SIF-1 involved in hydrogen bonds (left) and salt bridges (right), as identified from the crystal structure. **(B)** Evolutionary conservation profile of SIF-1 generated with ConSurf using SSLP F-box proteins from *C. tropicalis* as homologs. The THOR-1-binding surface of SIF-1, including the structural element encompassing residues 100–102, is poorly conserved among SIF-1-related proteins, whereas the opposite surface is highly conserved.

**Figure S7. Mutations in the THOR-1–SIF-1 interface result in lethality.** Quantification of embryonic lethality among the progeny of animals heterozygous for the SIF-1 Arg^142^Glu mutation (right). The ∼25% embryonic lethality is consistent with a homozygous-lethal phenotype. Genotypes of surviving progeny from four SIF-1 mutant lines carrying different combinations of substitutions at the key THOR-1-interacting residues Arg^142^, Thr^147^, and Arg^151^. Only homozygous wild-type and heterozygous mutant animals were recovered and were fertile; no homozygous mutant animals were obtained, indicating that these residues are essential for SIF-1 antidote function (left).

**Figure S8. Sequence alignment and pairwise amino-acid identity of SLOW toxins and their linked GROW antidotes. (A)** The alignment includes the toxins SLOW-1, SLOW-2, and SLOW-3 and **(B)** their corresponding antidotes, GROW-1, GROW-2, and GROW-3.

## Supplementary Tables

**Table S1.** Candidate genes for the EG6180 Chr. V TA.

**Table S2.** Candidate genes for the NIC203 Chr. V TA.

**Table S3.** The family of SSLP F-box proteins.

**Table S4**. AlphaFold 3 predictions of interactions between the SGYN-1 antidote and *C. tropicalis* SKR proteins.

**Table S5**. X-ray diffraction data collection and crystallographic refinement statistics for the THOR-1–SIF-1 complex. Values in parentheses correspond to the highest-resolution shell.

**Table S6**. Presence and absence of toxin–antidote elements across *C. tropicalis* natural isolates.

**Table S7**. Pairs of *C. tropicalis* isolates carrying opposing toxin–antidote haplotypes collected at identical geographic coordinates.

**Table S8**. List of all the strains used in this study

**Table S9**. Summary of raw phenotyping and genotyping data for all genetic crosses.

**Table S10**. Summary of all primers and guide RNAs, and HDR donor templates.

## Supplementary Data

**Data S1.** Interactive map showing the geographic distribution of *C. tropicalis* natural isolates and their toxin–antidote haplotypes.

