## Supplementary Information for "Conflicts and coalitions among selfish toxin-antidote elements shape the evolution of hyperdivergent haplotypes"

### **Supplemental Information**

Supplemental Figures S1-S8
Supplemental Tables S1-S11
Data S1

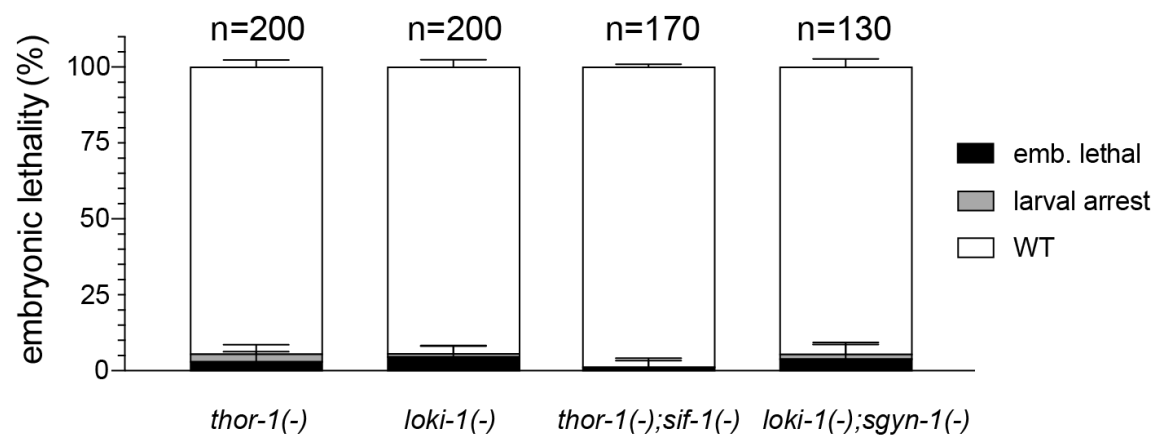

**Figure S1. Phenotypic characterization of null mutant lines.** Neither single-toxin mutants nor double toxin–antidote mutant combinations exhibited developmental defects above background levels. These results indicate that the corresponding genes are dispensable for normal *C. tropicalis* embryonic development, consistent with their proposed roles as selfish genetic elements.

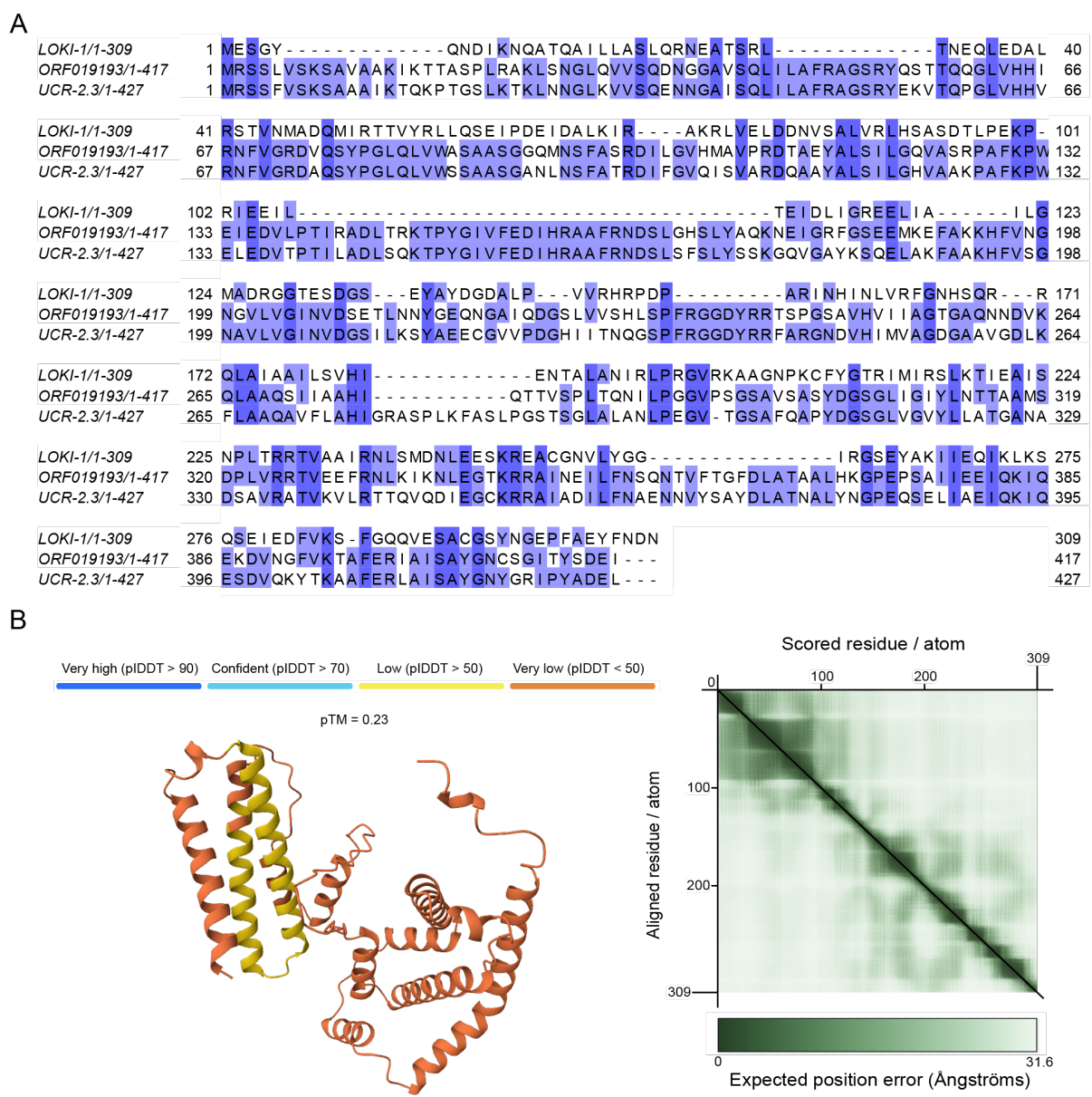

**Figure S2. The *loki-1* toxin is homologous to *C. elegans ucr-2.3* but is highly divergent. (A)** Protein sequence alignment of *C. tropicalis* LOKI-1, the *C. tropicalis* *ucr-2.3* ortholog ORF019193, and *C. elegans* UCR-2.3. **(B)** AlphaFold3 prediction of the LOKI-1 protein structure. The overall confidence of the predicted model is very low, providing limited insight into the potential mechanism of LOKI-1 toxicity.

A

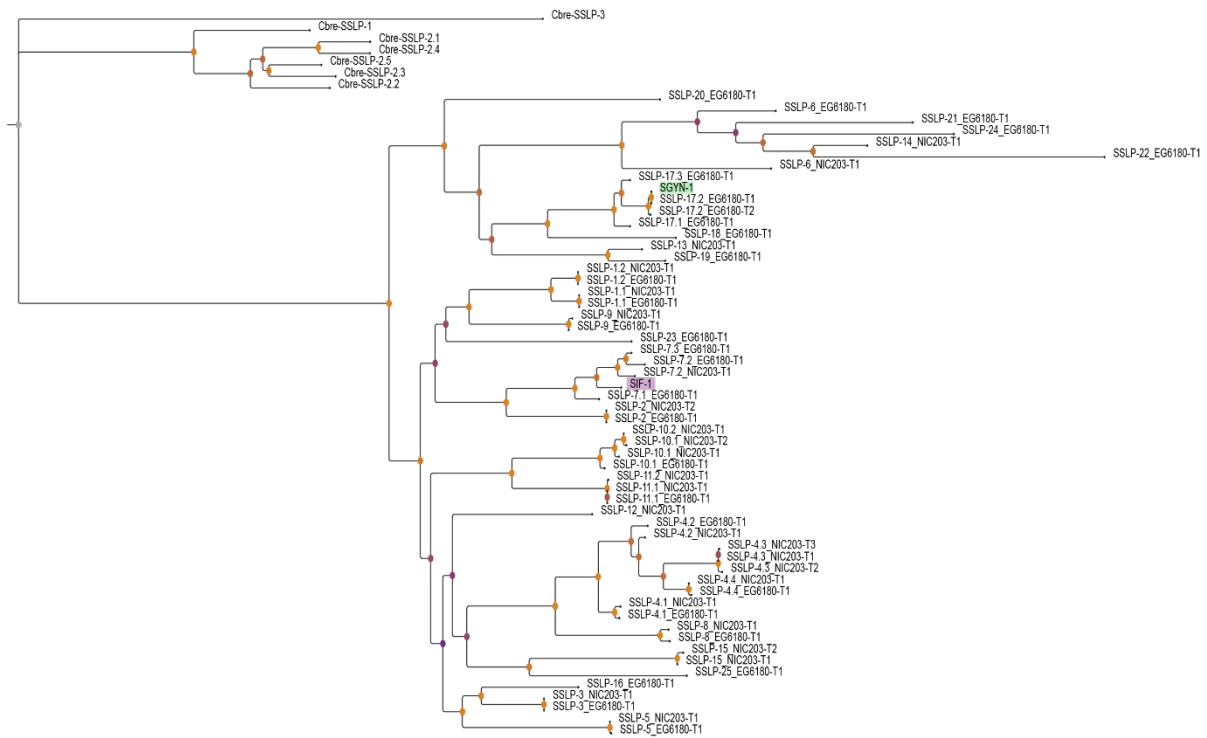

B

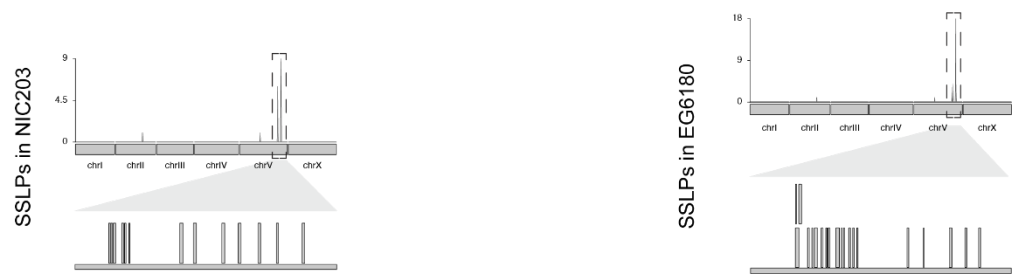

**Figure S3. Phylogenetic relationships among SGYN-1, SIF-1, and the SSLP family of F-box proteins and their genomic distribution. (A)** Phylogenetic tree including all annotated SSLP (SIF- and SGYN-like protein) F-box proteins from the *C. tropicalis* wild isolates NIC203 and EG6180 and from *C. brenneri*, together with the antidotes SGYN-1 and SIF-1. **(B)** Genomic distribution of SSLP genes in NIC203 (right) and EG6180 (left). In both isolates, SSLP genes form clusters on the right arm of chromosome V.

### A THOR-1

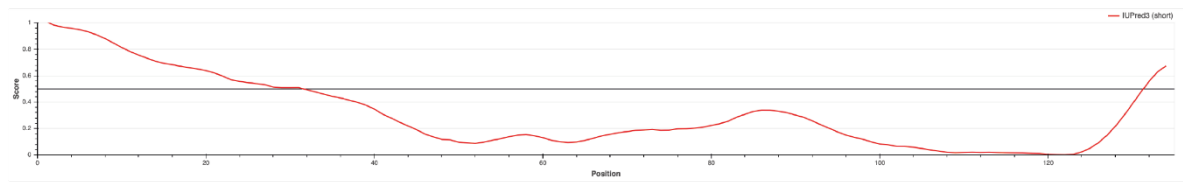

### SIF-1

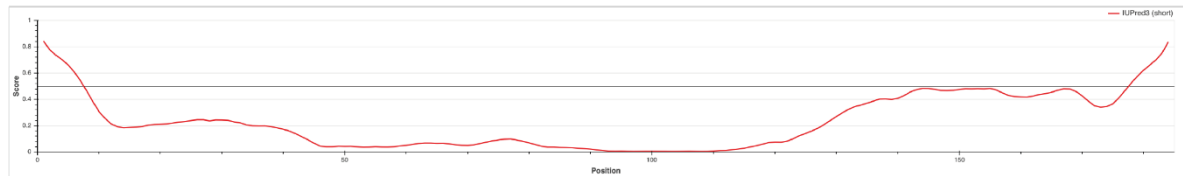

## B

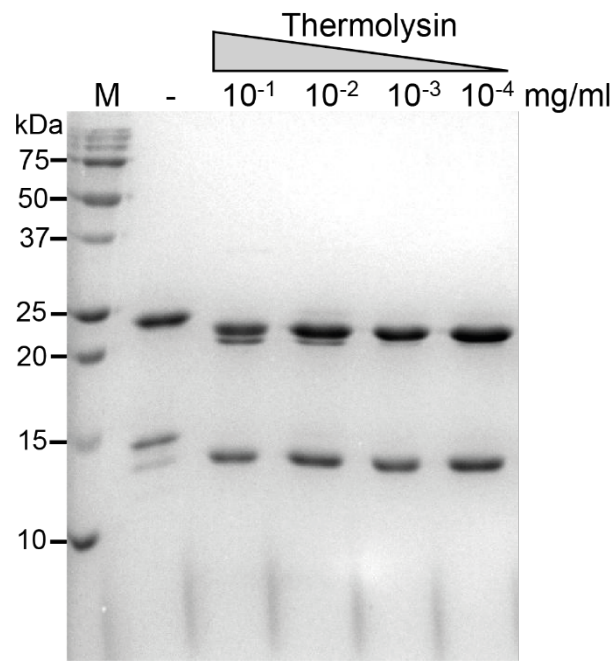

**Figure S4. Flexible regions in THOR-1 and SIF-1 and limited proteolysis of the THOR-1–SIF-1 complex.** (A) Predicted intrinsically disordered regions in THOR-1 (top) and SIF-1 (bottom), generated using IUPred2. (B) Limited proteolysis of the THOR-1–SIF-1 complex with increasing concentrations of thermolysin. Reproducible stable proteolytic fragments were observed. Proteins were visualized by Coomassie staining.

A

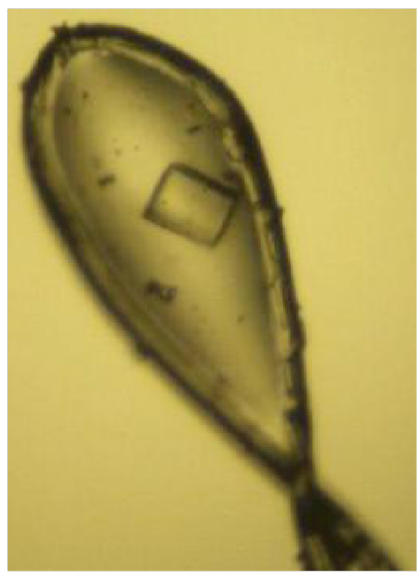

| P 21 21 2 | Compl. | Res. | Rmerge |
| --- | --- | --- | --- |
| Overall | 99.6% | 97.7-2.2 | 12.3 |
| Inner | 99.6% | 97.7-9.0 | 4.6 |
| Outer | 96.6% | 2.25-2.18 | 114.2 |

| a | b | c |
| --- | --- | --- |
| 89.97 Å | 97.72 Å | 40.32 Å |
| $\alpha$ | $\beta$ | $\gamma$ |
| 90 ° | 90 ° | 90 ° |

B

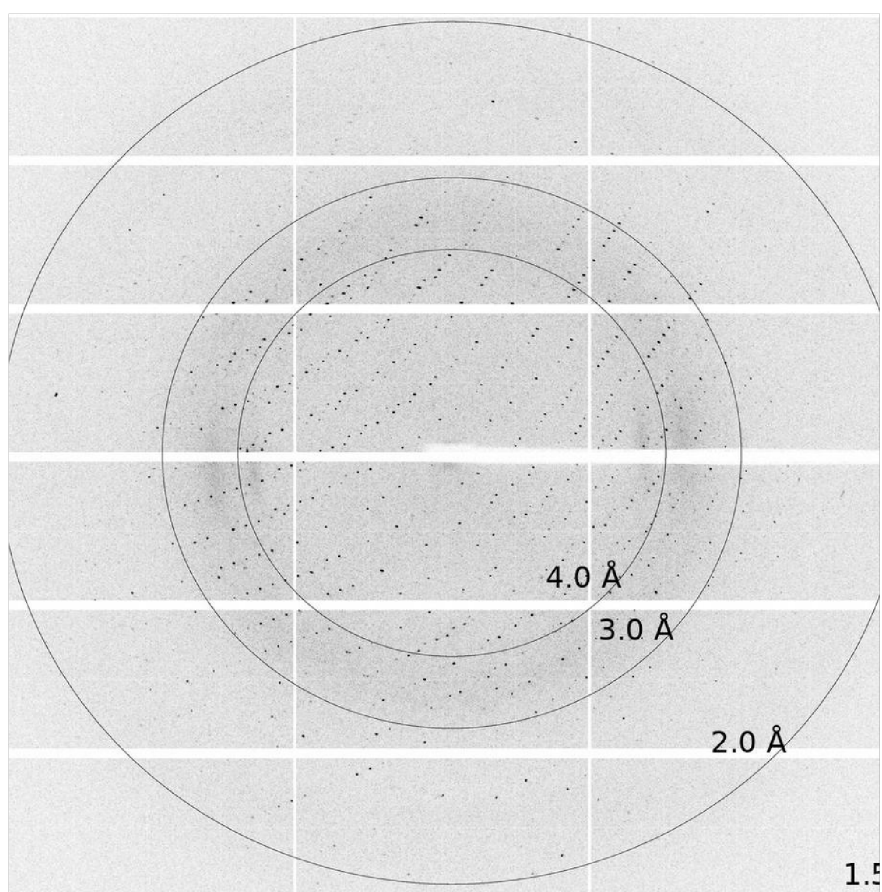

**Figure S5. Crystallization and X-ray diffraction of the THOR-1-SIF-1 complex.** (A) Representative crystal of the THOR-1-SIF-1 complex (left). Crystallographic data collection statistics (right). Space group, unit-cell parameters, completeness, resolution range, and Rmerge are shown for the THOR-1-SIF-1 crystal dataset. Inner- and outer-shell statistics correspond to 97.7–9.0 Å and 2.25–2.18 Å, respectively. (B) Representative X-ray diffraction image showing Bragg reflections extending to approximately 2.2 Å resolution.

A

| Hydrogen bonds |  |  |  | Salt bridges |  |  |  |
| --- | --- | --- | --- | --- | --- | --- | --- |
| ## | THOR-1 | Dist. [Å] | SIF-1 | ## | THOR-1 | Dist. [Å] | SIF-1 |
| 1 | B:PHE 34[ N ] | 2.83 | A:TYR 102[ O ] | 1 | B:ARG 35[ NE ] | 3.92 | A:ASP 69[ OD1] |
| 2 | B:ARG 35[ NH1] | 2.95 | A:ASP 73[ OD1] | 2 | B:ARG 35[ NE ] | 3.85 | A:ASP 69[ OD2] |
| 3 | B:ARG 35[ NH2] | 3.33 | A:ASP 73[ OD1] | 3 | B:ARG 35[ NH1] | 2.95 | A:ASP 73[ OD1] |
| 4 | B:ARG 35[ NH2] | 3.14 | A:ASP 73[ OD2] | 4 | B:ARG 35[ NH1] | 3.98 | A:ASP 73[ OD2] |
| 5 | B:ALA 36[ N ] | 2.99 | A:TRP 100[ O ] | 5 | B:ARG 35[ NH2] | 3.33 | A:ASP 73[ OD1] |
| 6 | B:ARG 73[ NH2] | 3.73 | A:GLU 136[ OE1] | 6 | B:ARG 35[ NH2] | 3.14 | A:ASP 73[ OD2] |
| 7 | B:THR 118[ OG1] | 3.21 | A:GLU 138[ OE1] | 7 | B:LYS 66[ NZ ] | 3.27 | A:ASP 73[ OD2] |
| 8 | B:ILE 124[ N ] | 3.49 | A:TYR 102[ OH ] | 8 | B:ARG 73[ NH2] | 3.73 | A:GLU 136[ OE1] |
| 9 | B:GLU 28[ O ] | 3.09 | A:LYS 106[ NZ ] | 9 | B:GLU 30[ OE1] | 3.17 | A:LYS 106[ NZ ] |
| 10 | B:GLU 30[ OE2] | 2.77 | A:LYS 106[ NZ ] | 10 | B:GLU 30[ OE2] | 2.77 | A:LYS 106[ NZ ] |
| 11 | B:PHE 34[ O ] | 2.79 | A:TYR 102[ N ] | 11 | B:GLU 84[ OE1] | 3.31 | A:ARG 151[ NH1] |
| 12 | B:GLY 80[ O ] | 3.57 | A:ARG 134[ NH2] | 12 | B:GLU 84[ OE1] | 2.96 | A:ARG 151[ NH2] |
| 13 | B:PRO 82[ O ] | 2.92 | A:ARG 151[ NH1] | 13 | B:GLU 84[ OE2] | 2.98 | A:ARG 151[ NH1] |
| 14 | B:GLU 84[ OE1] | 2.96 | A:ARG 151[ NH2] | 14 | B:GLU 84[ OE2] | 3.69 | A:ARG 151[ NH2] |
| 15 | B:GLU 84[ OE2] | 2.98 | A:ARG 151[ NH1] | 15 | B:ASP 121[ OD1] | 2.72 | A:ARG 142[ NE ] |
| 16 | B:THR 118[ O ] | 3.79 | A:ARG 151[ NH2] | 16 | B:ASP 121[ OD1] | 2.71 | A:ARG 142[ NH2] |
| 17 | B:THR 118[ OG1] | 3.38 | A:ARG 151[ NH2] | 17 | B:ASP 121[ OD2] | 3.76 | A:ARG 142[ NE ] |
| 18 | B:ASP 121[ OD1] | 2.72 | A:ARG 142[ NE ] |  |  |  |  |
| 19 | B:ASP 121[ OD1] | 2.71 | A:ARG 142[ NH2] |  |  |  |  |
| 20 | B:ASP 121[ OD2] | 2.76 | A:THR 147[ OG1] |  |  |  |  |

B

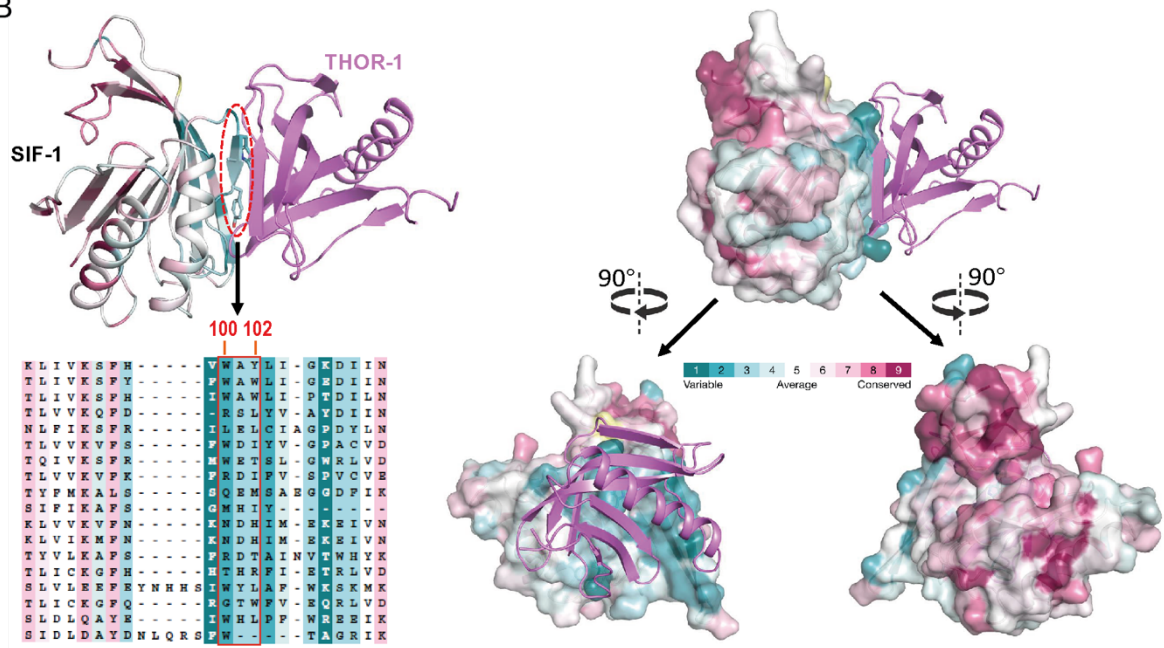

**Figure S6. Structural details of the THOR-1–SIF-1 interaction interface.** (A) Residues in THOR-1 and SIF-1 involved in hydrogen bonds (left) and salt bridges (right), as identified from the crystal structure. (B) Evolutionary conservation profile of SIF-1 generated with ConSurf using SSFP F-box proteins from *C. tropicalis* as homologs. The THOR-1-binding surface of SIF-1, including the structural element encompassing residues 100–102, is poorly conserved among SIF-1-related proteins, whereas the opposite surface is highly conserved.

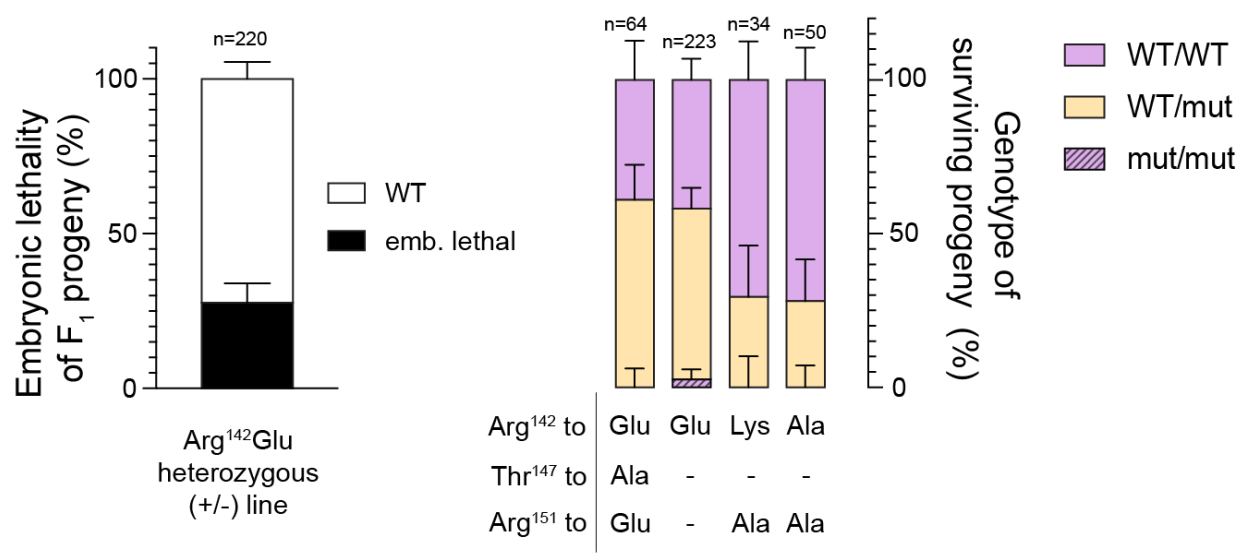

**Figure S7. Mutations in the THOR-1-SIF-1 interface result in lethality.** Quantification of embryonic lethality among the progeny of animals heterozygous for the SIF-1 Arg<sup>142</sup>Glu mutation (left). The ~25% embryonic lethality is consistent with a homozygous-lethal phenotype. Genotypes of surviving progeny from four SIF-1 mutant lines carrying different combinations of substitutions at the key THOR-1-interacting residues Arg<sup>142</sup>, Thr<sup>147</sup>, and Arg<sup>151</sup>. Only homozygous wild-type and heterozygous mutant animals were recovered and were fertile; no homozygous mutant animals were obtained, indicating that these residues are essential for SIF-1 antidote function (right).

A

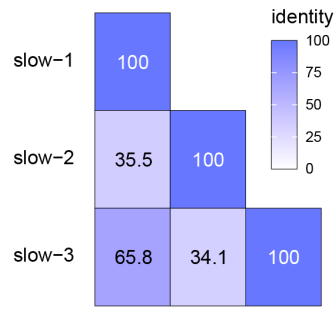

|  |  |  |  |  |  |  |  |  |  |  |  |  |  |  |  |  |  |  |  |
| --- | --- | --- | --- | --- | --- | --- | --- | --- | --- | --- | --- | --- | --- | --- | --- | --- | --- | --- | --- |
| slow-2/1-357 | 1 | MELTPRV | SCELSK | LATVYE | AHORIL | LTVSSQ | SEEEV | VVGEVE | QSLQ | ELNV | SHCH | KKFE | LENVI | 61 |  |  |  |  |  |
| slow-1/1-364 | 1 | MELV-- | TLCDLSKLA | ILHET | LQKVLDV | SCQTEEE | IKKEE | GVLR | EITAA | ADCKQ | MESLERIA |  |  | 59 |  |  |  |  |  |
| slow-3/1-371 | 1 | MELV-- | TLCDLSKLA | ILHEN | LQNVLDV | SCQSESE | LFKKKE | INLQ | LTGA | DDQM | QLMLERIA |  |  | 59 |  |  |  |  |  |
| slow-2/1-357 | 62 | LKSW | LEFRR | IDEIA | APDR | TLML | LOYRRA | KWIL | LDHL | FETSR | NEKE | CSKK | RVLV | FYGY | FFDP | 122 |  |  |  |
| slow-1/1-364 | 60 | FNSF | VTSD | FRFNE | LPELD | KETLI | KQYRT | SMMAL | NRLLE | GSKY | EREQ | SEKLV | VRRR | FYTK | 120 |  |  |  |  |
| slow-3/1-371 | 60 | FNSF | VTSD | FRFNE | LPELD | KETLI | KQYRT | SMMAL | NRLLE | GSKY | EREQ | SEKLV | VRRR | FYTK | 120 |  |  |  |  |
| slow-2/1-357 | 123 | QMPF | LLID | SNPR | SVDA | LEDV | NKRR | LNQM | WDIV | SAFI | ENKVT | VEE | GYM | HLSL | FWKP | Q-- | N | 180 |  |
| slow-1/1-364 | 121 | KLKP | H-- | EET | SENS | VAREE | EHKK | RVD | SVWE | VNVG | I | EANVT | SEEI | AALH | MLFW | ESSD | PS | 179 |  |
| slow-3/1-371 | 121 | ELKP | N-- | DEET | ENCV | ARK | EVHQ | KKAD | SVWE | VVGAI | A | EANP | SSEE | I | VALH | MLFW | NSPR | D | 178 |
| slow-2/1-357 | 181 | ESCL | SLECO | KIME | ERRNS | TEKVI | ENCWR | KEGCP | EGRK | DVITE | EQMS | VSA | LAH | SSSV | F | SNVQ | 241 |  |  |
| slow-1/1-364 | 180 | ETSL | SKSQ | QOKI | LKER | QAKI | ITMS | LEKY | HESK | GINAN | RKEV | IKKK | LKIS | IGT | LVHQ | ITDD | FRLG | 240 |  |
| slow-3/1-371 | 179 | EMNL | SENG | RRILE | ERQKS | IMTLD | KLYE | AKGI | VSI | R | IEVI | KNL | KIL | LIEK | LVNG | ITDA | FRLD | 239 |  |
| slow-2/1-357 | 242 | NSHP | MISS | DFLDS | DFDGI | KSTL | RETSP | DD----- | VPEV | PAVT | ----- | ----- | ----- | ----- | ----- | ----- | ----- | 284 |  |
| slow-1/1-364 | 241 | NRKT | FESS | -FINS | IDFGL | DET | FSKR | SDDEN | LPVVA | LPVAA | PAF | ----- | ----- | ----- | ----- | ----- | ----- | 288 |  |
| slow-3/1-371 | 240 | NQKM | YSS- | -FINS | IDFGL | DET | LSVVA | DEH----- | VPAT | VPAP | PAST | VPAP | SPFD | NLS | NMN | 293 |  |  |  |
| slow-2/1-357 | 285 | LFGK | EWST | DRAS | TFSF | SGVLA | VLAAL | VLVLF | FTTE | KDKRT | TEFMI | LVIS | SFVAA | I | FSVW | LFYT | G | 345 |  |
| slow-1/1-364 | 289 | LFGK | KVNT | ARLGT | FLSG | IAIV | VGAV | STFAS | VDKD | HDVT | TEFVI | ILFV | FSLV | VGLI | LMAY | LPFAP | 349 |  |  |
| slow-3/1-371 | 294 | LFGK | KVNT | ARLGT | FLSG | VVVV | I | GA | VSSYA | SVDK | EHVT | TEFVI | ILFV | LT | LVVG | LTMT | VPFAP | 354 |  |
| slow-2/1-357 | 346 | TKR | EFK | KHFVK | - - - - |  |  |  |  |  |  |  |  |  |  |  |  | 357 |  |
| slow-1/1-364 | 350 | HA | ILCLR | RFVGD | WFG- | - - |  |  |  |  |  |  |  |  |  |  |  | 364 |  |
| slow-3/1-371 | 355 | HI | ISFL | QRFM | GP | DGSQ | I |  |  |  |  |  |  |  |  |  |  | 371 |  |

B

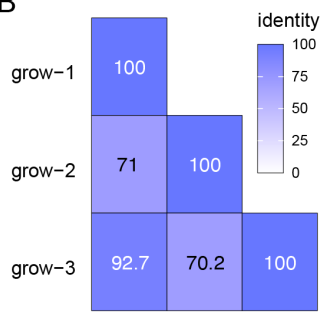

|  |  |  |  |  |  |  |  |  |  |  |  |  |  |  |  |  |  |  |  |  |  |  |
| --- | --- | --- | --- | --- | --- | --- | --- | --- | --- | --- | --- | --- | --- | --- | --- | --- | --- | --- | --- | --- | --- | --- |
| grow-2/1-124 | 1 | MGNNAEV | ORLKEINS | IKOTTALY | IKILKGI | AEYDQI | IDS | KLVNV | KGDFEL | YHFSRD | 60 |  |  |  |  |  |  |  |  |  |  |  |
| grow-1/1-124 | 1 | MGNDAKA | IRRLTKA | ELIEQVT | ADRMAYLKAC | IEYDQI | IDS | KLVN | KGDFEL | YHFSRD | 60 |  |  |  |  |  |  |  |  |  |  |  |
| grow-3/1-124 | 1 | MGNDAAET | IRRLTKA | IFMEQI | IERMAYLKAC | IEYDQI | IDS | KLVN | KGDFEL | YHFSRD | 60 |  |  |  |  |  |  |  |  |  |  |  |
| grow-2/1-124 | 61 | KKCLA | EF | DHLP | SDEE | KAICYKV | KDQFKA | IEEA | ERRR | EKS | IFVNR | VSR | SI | WT | PI | K | SFF | NRRH | 120 |  |  |  |
| grow-1/1-124 | 61 | KKCLA | EF | DHLP | S | NKKRRVD | SWLL | I | NNQ | FKA | IEEA | ERRR | EKS | IFVNR | VSR | SI | WT | PI | K | SFF | NRRH | 120 |
| grow-3/1-124 | 61 | KKCLA | EF | DHLP | S | NKKRRVD | SWLL | I | NNQ | FKA | IEEA | ERRR | EKS | IFVNR | VSR | SI | WT | PI | K | SFF | NRRH | 120 |
| grow-2/1-124 | 121 | L | PQK |  |  |  |  |  |  |  |  |  |  |  |  |  |  |  |  |  | 124 |  |
| grow-1/1-124 | 121 | L | PQK |  |  |  |  |  |  |  |  |  |  |  |  |  |  |  |  |  | 124 |  |
| grow-3/1-124 | 121 | L | PQK |  |  |  |  |  |  |  |  |  |  |  |  |  |  |  |  |  | 124 |  |

**Figure S8. Sequence alignment and pairwise amino-acid identity of SLOW toxins and their linked GROW antidotes.** (A) The alignment includes the toxins SLOW-1, SLOW-2, and SLOW-3 and (B) their corresponding antidotes, GROW-1, GROW-2, and GROW-3.

**Table S1. Candidate genes for the EG6180 Chr. V TA.**

| **ORF_ID**<br>**EG6180** | **NIC203\|evalue\|query_cov\|p**<br>**erc_ident** | **protein_ID_C.elegans\|evalue\|query_co**<br>**v\|perc_ident** | **TPM** |
| --- | --- | --- | --- |
| FUN_019<br>184 | FUN_019751\|0.0\|100\|100.00<br>0 | tr\|O18693\|O18693_CAEEL\|0.0\|100\|93.52<br>8 | 189.4171<br>60 |
| FUN_019<br>185 | FUN_019752\|0.0\|100\|97.323 | tr\|G5EG72\|G5EG72_CAEEL\|0.0\|100\|92.<br>939 | 1.398273 |
| FUN_019<br>186 | FUN_019753\|2.56e-171\|100\|100.000 | tr\|O17824\|O17824_CAEEL\|1.68e-127\|100\|76.271 | 3.587802 |
| FUN_019<br>187 | FUN_019754\|0.0\|100\|99.803 | sp\|O17828\|PRDE1_CAEEL\|0.0\|83\|57.00<br>9 | 9.860572 |
| FUN_019<br>188 | FUN_019755\|1.40e-174\|97\|100.000 | tr\|O17827\|O17827_CAEEL\|1.63e-133\|100\|77.459 | 12.89370<br>6 |
| FUN_019<br>189 | FUN_019756\|0.0\|100\|99.383 | tr\|H2FLJ4\|H2FLJ4_CAEEL\|0.0\|94\|86.334 | 24.09944<br>5 |
| FUN_019<br>190 | FUN_019757\|3.38e-35\|46\|92.982 | tr\|H2FLJ5\|H2FLJ5_CAEEL\|1.60e-33\|42\|92.308 | 1.863198 |
| FUN_019<br>191 | FUN_019759\|8.19e-57\|100\|91.463 | tr\|O62115\|O62115_CAEEL\|3.70e-19\|94\|48.718 | 0.245289 |
| FUN_019<br>192 | FUN_019760\|1.71e-53\|100\|91.463 | tr\|O62114\|O62114_CAEEL\|8.63e-24\|91\|53.333 | 0.180248 |
| FUN_019<br>193 | FUN_019761\|0.0\|100\|96.643 | tr\|Q9TZ33\|Q9TZ33_CAEEL\|0.0\|100\|59.8<br>13 | 70.84793<br>1 |
| **loki-1** | **FUN_019761\|3.10e-**<br>**17\|47\|29.885** | **tr\|Q9TZ33\|Q9TZ33_CAEEL\|9.10e-**<br>**05\|68\|27.007** | **22.25400**<br>**2** |
| **sgyn-1** | **sslp-13\|6.16e-82\|92\|54.936** | **tr\|O62012\|O62012_CAEEL\|2.04e-**<br>**12\|64\|31.624** | **17,78772**<br>**4** |
| sslp-17.1 | sslp-13\|3.98e-80\|95\|55.507 | tr\|O62012\|O62012_CAEEL\|4.87e-13\|51\|30.769 | 0.302929 |
| sslp-17.2 | sslp-13\|6.27e-66\|94\|51.442 | tr\|O62012\|O62012_CAEEL\|3.25e-08\|67\|28.846 | 8.966325 |
| sslp-17.3 | sslp-13\|1.05e-81\|98\|51.779 | tr\|O62012\|O62012_CAEEL\|4.04e-12\|48\|29.915 | 10.18763<br>4 |
| FUN_019<br>200 | sslp-11.2\|2.05e-17\|79\|54.237 | tr\|A0A4V0ILL1\|A0A4V0ILL1_CAEEL\|1.68<br>e-05\|40\|52.381 | 1.528112 |
| sslp-18 | sslp-13\|4.56e-73\|92\|50.000 | tr\|O62012\|O62012_CAEEL\|5.46e-12\|47\|29.915 | 3.632865 |
| sslp-19 | sslp-13\|3.82e-91\|96\|72.472 | tr\|G5EGT7\|G5EGT7_CAEEL\|9.46e-05\|40\|42.500 | 0.248699 |
| sslp-20 | FUN_019762\|5.36e-45\|42\|79.012 | tr\|Q19680\|Q19680_CAEEL\|5.74e-04\|23\|35.484 | 3.883801 |
| sslp-21 | sslp-14\|7.49e-88\|87\|57.872 | tr\|Q9U2X6\|Q9U2X6_CAEEL\|1.75e-12\|43\|33.071 | 0.568437 |
| FUN_019<br>205 | sslp-14\|2.38e-56\|83\|86.957 | tr\|Q9U2X6\|Q9U2X6_CAEEL\|3.77e-13\|69\|35.897 | 0.681304 |
| sslp-6 | sslp-6\|2.08e-134\|100\|82.063 | tr\|P90770\|P90770_CAEEL\|1.59e-04\|52\|26.437 | 0.235955 |
| sslp-22 | sslp-14\|2.91e-85\|95\|55.752 | tr\|Q9U2X6\|Q9U2X6_CAEEL\|8.71e-14\|48\|31.008 | 4.996680 |
| FUN_019<br>208 | FUN_019766\|0.0\|91\|97.122 | tr\|O17826\|O17826_CAEEL\|1.02e-141\|98\|61.056 | 1.137449 |
| sslp-23 | sslp-9\|1.26e-71\|99\|45.726 | tr\|Q9TZ26\|Q9TZ26_CAEEL\|7.68e-10\|38\|26.596 | 12.77702<br>7 |
| sslp-24 | sslp-14\|7.60e-67\|75\|61.111 | tr\|P91038\|P91038_CAEEL\|9.51e-05\|54\|23.729 | 0.761869 |
| sslp-7.1 | sif-1\|5.96e-109\|95\|78.212 | tr\|Q966H8\|Q966H8_CAEEL\|9.71e-05\|51\|25.926 | 1.047432 |
| sslp-7.2 | sslp-7.2\|1.84e-114\|99\|84.066 | sp\|O18086\|NH218_CAEEL\|1.06e-04\|53\|30.435 | 2.874153 |
| sslp-25 | sslp-2\|2.01e-50\|92\|45.405 | tr\|Q9N5X0\|Q9N5X0_CAEEL\|2.25e-05\|24\|29.787 | 0.000000 |
| sslp-7.3 | sslp-7.2\|6.45e-126\|99\|89.011 | tr\|A0A1T5HUI7\|A0A1T5HUI7_CAEEL\|4.45e-04\|31\|32.692 | 2.470244 |
| FUN_019215 | thor-1\|1.68e-26\|88\|35.833 | tr\|O16550\|O16550_CAEEL\|1.17e-04\|48\|26.471 | 33.403049 |
| FUN_019216 | FUN_019771\|4.80e-49\|100\|100.000 | sp\|O17570\|RL38_CAEEL\|4.54e-48\|100\|98.571 | 1824.409424 |
| FUN_019217 | FUN_019772\|0.0\|99\|83.943 | tr\|G5EE87\|G5EE87_CAEEL\|0.0\|99\|97.059 | 9.373868 |
| FUN_019219 | FUN_019774\|0.0\|100\|98.701 | tr\|O17829\|O17829_CAEEL\|0.0\|100\|86.774 | 7.010653 |
| FUN_019220 | FUN_005301\|3.73e-05\|74\|26.027 | tr\|G5EE87\|G5EE87_CAEEL\|1.05e-40\|91\|60.000 | 5.434507 |
| sslp-8 | sslp-8\|1.90e-147\|100\|93.204 | tr\|G5EEP4\|G5EEP4_CAEEL\|7.27e-10\|52\|33.333 | 0.000000 |
| FUN_019222 | FUN_019777\|0.0\|99\|95.278 | sp\|Q09336\|FB224_CAEEL\|2.78e-27\|80\|24.786 | 1.248814 |
| sslp-1.2 | sslp-1.2\|2.20e-104\|100\|100.000 | tr\|Q9N4E1\|Q9N4E1_CAEEL\|0.002\|34\|36.170 | 0.365764 |
| FUN_019224 | FUN_022348\|6.18e-05\|55\|61.538 | tr\|Q21948\|Q21948_CAEEL\|2.93e-04\|30\|52.174 | 34.414581 |
| FUN_019225 | FUN_019781\|1.22e-52\|69\|97.297 | sp\|O17570\|RL38_CAEEL\|6.35e-12\|29\|75.758 | 58.752430 |
| sslp-9 | sslp-9\|5.93e-178\|100\|98.739 | tr\|Q9U2X6\|Q9U2X6_CAEEL\|5.75e-12\|43\|33.058 | 1.511286 |
| FUN_019227 | FUN_019783\|1.06e-76\|100\|100.000 | tr\|Q9XX88\|Q9XX88_CAEEL\|1.14e-06\|95\|28.571 | 0.000000 |
| FUN_019228 | FUN_019784\|1.58e-39\|100\|93.333 | tr\|O62012\|O62012_CAEEL\|1.63e-12\|93\|40.323 | 0.000000 |
| FUN_019229 | FUN_019785\|3.94e-145\|97\|97.949 | tr\|Q966E4\|Q966E4_CAEEL\|2.81e-12\|80\|26.347 | 0.000000 |
| FUN_019230 | FUN_019786\|1.43e-66\|100\|100.000 | tr\|G5EFR0\|G5EFR0_CAEEL\|3.19e-04\|44\|34.091 | 0.000000 |
| sslp-10.1 | sslp-10.1\|1.45e-147\|87\|99.492 | tr\|Q9TZ24\|Q9TZ24_CAEEL\|1.39e-11\|49\|31.915 | 0.909048 |
| FUN_019232 | FUN_019798\|5.72e-59\|100\|100.000 | tr\|Q9N5W8\|Q9N5W8_CAEEL\|2.50e-07\|99\|30.909 | 0.000000 |
| FUN_019233 | FUN_019794\|0.0\|100\|99.149 | tr\|O62012\|O62012_CAEEL\|1.89e-28\|51\|45.082 | 0.288387 |
| sslp-11.1 | sslp-11.1\|3.47e-175\|100\|100.000 | tr\|Q9U2X6\|Q9U2X6_CAEEL\|1.11e-10\|48\|28.030 | 0.046284 |
| FUN_019235 | FUN_019803\|2.63e-69\|100\|100.000 | tr\|P91468\|P91468_CAEEL\|7.99e-04\|47\|31.111 | 7.883569 |

**Table S2. Candidate genes for the NIC203 Chr. V TA.**

| **ORF_ID**<br>**NIC203** | **EG6180\|evalue\|query_cov\|p**<br>**erc_ident** | **protein_ID_C.elegans\|evalue\|query_co**<br>**v\|perc_ident** | **TPM** |
| --- | --- | --- | --- |
| FUN_019<br>751 | FUN_019184\|0.0\|100\|100.000 | tr\|O18693\|O18693_CAEEL\|0.0\|100\|93.52<br>8 | 484.574<br>371 |
| FUN_019<br>752 | FUN_019185\|0.0\|100\|97.323 | tr\|G5EG72\|G5EG72_CAEEL\|0.0\|100\|95.4<br>90 | 1.67268<br>9 |
| FUN_019<br>753 | FUN_019186\|6.07e-171\|100\|100.000 | tr\|O17824\|O17824_CAEEL\|1.68e-127\|100\|76.271 | 2.21616<br>5 |
| FUN_019<br>754 | FUN_019187\|0.0\|100\|99.803 | sp\|O17828\|PRDE1_CAEEL\|0.0\|83\|57.00<br>9 | 8.10036<br>7 |
| FUN_019<br>755 | FUN_019188\|3.26e-174\|98\|100.000 | tr\|O17827\|O17827_CAEEL\|7.62e-129\|98\|77.542 | 5.95403 |
| FUN_019<br>756 | FUN_019189\|0.0\|100\|99.383 | tr\|H2FLJ4\|H2FLJ4_CAEEL\|0.0\|94\|85.900 | 29.9100<br>57 |
| FUN_019<br>757 | FUN_017669\|1.63e-66\|75\|54.598 | sp\|P92001\|SRD18_CAEEL\|2.05e-45\|72\|46.429 | 1.12761<br>4 |
| FUN_019<br>758 | FUN_017669\|9.41e-43\|100\|66.304 | tr\|Q9UAX5\|Q9UAX5_CAEEL\|2.99e-32\|95\|52.874 | 0.09158<br>5 |
| FUN_019<br>759 | FUN_019191\|1.94e-56\|100\|91.463 | tr\|O62165\|O62165_CAEEL\|2.16e-21\|94\|50.649 | 0 |
| FUN_019<br>760 | FUN_019192\|4.05e-53\|100\|91.463 | tr\|O62114\|O62114_CAEEL\|4.94e-25\|91\|56.000 | 0 |
| FUN_019<br>761 | FUN_019193\|0.0\|100\|96.643 | tr\|Q9TZ33\|Q9TZ33_CAEEL\|0.0\|100\|60.7<br>48 | 61.0598<br>91 |
| FUN_019<br>762 | sslp-18\|2.96e-55\|97\|53.793 | tr\|Q9XX86\|Q9XX86_CAEEL\|1.37e-10\|48\|51.220 | 4.40386<br>9 |
| sslp-13 | sslp-19\|1.22e-90\|67\|72.472 | tr\|Q9TZ24\|Q9TZ24_CAEEL\|2.31e-11\|31\|32.468 | 3.64097<br>3 |
| sslp-14 | sslp-21\|1.51e-87\|98\|57.872 | tr\|H2KZR8\|H2KZR8_CAEEL\|8.05e-16\|60\|30.201 | 6.13824<br>6 |
| sslp-6 | sslp-6\|5.94e-134\|78\|82.063 | tr\|H2KZR8\|H2KZR8_CAEEL\|1.81e-11\|62\|26.087 | 3.71188<br>9 |
| FUN_019<br>766 | FUN_019208\|0.0\|92\|97.122 | tr\|O17826\|O17826_CAEEL\|7.62e-145\|98\|62.416 | 0.69011<br>9 |
| sslp-15 | sslp-2\|7.20e-68\|83\|49.537 | tr\|G5EEP4\|G5EEP4_CAEEL\|2.30e-07\|62\|25.414 | 4.58105<br>9 |
| sslp-7.2 | sslp-7.3\|1.53e-125\|99\|89.011 | tr\|Q9XTR6\|Q9XTR6_CAEEL\|2.14e-04\|45\|21.739 | 3.35983<br>6 |
| **sif-1** | **sslp-7.3\|2.17e-**<br>**110\|100\|80.874** | **sp\|P27715\|UNC13_CAEEL\|0.001\|56\|28.**<br>**571** | **45.8645**<br>**94** |
| **thor-1** | **FUN_019215\|3.92e-**<br>**26\|90\|35.833** | **tr\|Q9XU54\|Q9XU54_CAEEL\|2.38e-**<br>**04\|63\|26.437** | **19.6350**<br>**92** |
| FUN_019<br>771 | FUN_019216\|1.14e-48\|100\|100.000 | sp\|O17570\|RL38_CAEEL\|4.54e-48\|100\|98.571 | 960.854<br>736 |
| FUN_019<br>772 | FUN_019217\|0.0\|98\|83.943 | tr\|G5EE87\|G5EE87_CAEEL\|0.0\|99\|92.97<br>2 | 11.8381<br>92 |
| FUN_019<br>774 | FUN_019219\|0.0\|100\|98.701 | tr\|O17829\|O17829_CAEEL\|0.0\|100\|86.77<br>4 | 5.55195 |
| FUN_019<br>775 | FUN_013309\|2.91e-04\|85\|29.851 | sp\|P19826\|VINC_CAEEL\|1.70e-04\|66\|30.769 | 9.42863<br>4 |
| sslp-8 | sslp-8\|4.97e-147\|91\|93.204 | tr\|Q9BKX3\|Q9BKX3_CAEEL\|2.87e-10\|69\|24.390 | 4.35125<br>4 |
| FUN_019<br>777 | FUN_019222\|0.0\|100\|95.278 | sp\|Q09336\|FB224_CAEEL\|1.07e-24\|80\|25.309 | 10.4101<br>73 |
| sslp-1.2 | sslp-1.1\|3.42e-133\|100\|83.178 | tr\|Q9U2X6\|Q9U2X6_CAEEL\|1.71e-08\|45\|26.804 | 2.51479 |
| FUN_019<br>779 | FUN_018690\|2.50e-34\|66\|60.674 | tr\|G5ED36\|G5ED36_CAEEL\|7.92e-11\|61\|65.625 | 1.15399<br>9 |
| FUN_019<br>780 | FUN_019187\|1.36e-54\|100\|97.531 | sp\|O17828\|PRDE1_CAEEL\|1.30e-30\|100\|58.537 | 43.4971<br>35 |
| FUN_019<br>781 | FUN_019225\|3.61e-52\|56\|97.297 | tr\|A0A5S9MQQ5\|A0A5S9MQQ5_CAEEL\|3.40e-04\|31\|33.333 | 46.3818<br>28 |
| sslp-9 | sslp-9\|1.41e-177\|100\|98.739 | tr\|Q9U2X6\|Q9U2X6_CAEEL\|2.63e-12\|43\|32.787 | 13.4131<br>62 |
| FUN_019<br>783 | FUN_019227\|2.52e-76\|100\|100.000 | tr\|Q9XX88\|Q9XX88_CAEEL\|1.14e-06\|95\|28.571 | 0 |
| FUN_019<br>784 | FUN_019228\|3.74e-39\|100\|93.333 | tr\|O62012\|O62012_CAEEL\|1.43e-14\|88\|45.763 | 0.23007<br>9 |
| FUN_019<br>785 | FUN_019229\|1.22e-144\|74\|97.949 | tr\|Q9TYZ8\|Q9TYZ8_CAEEL\|1.18e-13\|54\|34.058 | 2.43137 |
| FUN_019<br>786 | FUN_019230\|3.39e-66\|100\|100.000 | tr\|G5EFR0\|G5EFR0_CAEEL\|3.19e-04\|44\|34.091 | 0.70524<br>9 |
| sslp-10.1 | sslp-10.1\|8.89e-146\|96\|99.487 | tr\|G5EDF8\|G5EDF8_CAEEL\|1.10e-06\|38\|34.177 | 17.2840<br>94 |
| FUN_019<br>788 | FUN_019230\|5.28e-49\|97\|100.000 | tr\|Q2WF60\|Q2WF60_CAEEL\|0.001\|99\|24.658 | 0.16975 |
| FUN_019<br>789 | FUN_019229\|2.11e-128\|98\|97.674 | tr\|Q966E6\|Q966E6_CAEEL\|8.17e-11\|70\|31.250 | 1.84058<br>4 |
| sslp-10.2 | sslp-10.1\|8.89e-146\|96\|99.487 | tr\|G5EDF8\|G5EDF8_CAEEL\|1.10e-06\|38\|34.177 | 14.4170<br>43 |
| FUN_019<br>791 | FUN_019230\|1.09e-36\|95\|94.737 | tr\|O45701\|O45701_CAEEL\|8.14e-04\|62\|35.714 | 0.05214<br>6 |
| FUN_019<br>792 | FUN_009147\|1.27e-06\|27\|28.947 | tr\|O44492\|O44492_CAEEL\|5.11e-136\|96\|64.552 | 0.67657<br>4 |
| sslp-11.1 | sslp-11.1\|8.23e-175\|100\|100.000 | tr\|Q9U2X6\|Q9U2X6_CAEEL\|1.11e-10\|48\|28.030 | 0.53443<br>1 |
| FUN_019<br>794 | FUN_019233\|2.19e-180\|100\|99.149 | tr\|O62012\|O62012_CAEEL\|5.66e-29\|51\|45.902 | 3.77465<br>1 |
| FUN_019<br>795 | FUN_019232\|1.36e-58\|100\|100.000 | tr\|Q9N5W8\|Q9N5W8_CAEEL\|2.50e-07\|99\|30.909 | 2.14474<br>6 |
| FUN_019<br>796 | FUN_018174\|4.60e-04\|37\|27.451 | tr\|G5ED87\|G5ED87_CAEEL\|4.42e-04\|57\|38.776 | 0 |
| FUN_019<br>797 | FUN_006892\|2.60e-05\|91\|31.746 | tr\|O45378\|O45378_CAEEL\|1.29e-05\|65\|32.727 | 0.32348<br>8 |
| FUN_019<br>798 | FUN_019232\|1.36e-58\|100\|100.000 | tr\|Q9N5W8\|Q9N5W8_CAEEL\|2.50e-07\|99\|30.909 | 2.15216<br>3 |
| FUN_019<br>799 | FUN_019233\|7.33e-74\|94\|97.000 | tr\|O62012\|O62012_CAEEL\|6.66e-26\|90\|50.526 | 1.69302<br>7 |
| sslp-11.2 | sslp-11.1\|3.77e-172\|95\|99.561 | tr\|Q9U2X6\|Q9U2X6_CAEEL\|1.52e-10\|48\|26.866 | 0.47785<br>4 |
| FUN_019<br>801 | FUN_021701\|7.28e-07\|73\|26.087 | tr\|O44492\|O44492_CAEEL\|5.16e-61\|95\|66.667 | 0.54145<br>1 |
| FUN_019<br>802 | FUN_021702\|9.93e-05\|57\|25.641 | tr\|O44492\|O44492_CAEEL\|4.83e-56\|100\|60.432 | 0.86819<br>8 |
| FUN_019<br>803 | FUN_019235\|6.23e-69\|100\|100.000 | tr\|P91468\|P91468_CAEEL\|7.99e-04\|47\|31.111 | 34.2443<br>89 |

**Table S3. The family of SSLP F-box proteins.**

| gene | old_name | species | isolate | chr | start | end |
| --- | --- | --- | --- | --- | --- | --- |
| Cbre-sslp-1_CFB2252 | CAL2028526.1 | Cbre | CFB2252 | OZ038406.1 | 5091004 | 5091857 |
| Cbre-sslp-2.1_CFB2252 | CAL2030684.1 | Cbre | CFB2252 | OZ038406.1 | 15236999 | 15237810 |
| Cbre-sslp-2.2_CFB2252 | CAL2030685.1 | Cbre | CFB2252 | OZ038406.1 | 15238410 | 15239249 |
| Cbre-sslp-2.3_CFB2252 | CAL2030686.1 | Cbre | CFB2252 | OZ038406.1 | 15239833 | 15240671 |
| Cbre-sslp-2.4_CFB2252 | CAL2030687.1 | Cbre | CFB2252 | OZ038406.1 | 15241624 | 15242426 |
| Cbre-sslp-2.5_CFB2252 | CAL2030688.1 | Cbre | CFB2252 | OZ038406.1 | 15242887 | 15244107 |
| Cbre-sslp-3_CFB2252 | CAL2047023.1 | Cbre | CFB2252 | OZ038410.1 | 17493510 | 17497439 |
| sslp-1.1_NIC203 | FUN_006758 | Ctro | NIC203 | chrII | 8509345 | 8512721 |
| sslp-2_NIC203 | FUN_017344 | Ctro | NIC203 | chrV | 6517978 | 6521226 |
| sslp-3_NIC203 | FUN_019286 | Ctro | NIC203 | chrV | 11977694 | 11979178 |
| sslp-4.1_NIC203 | FUN_019300 | Ctro | NIC203 | chrV | 12014032 | 12014907 |
| sslp-4.2_NIC203 | FUN_019301 | Ctro | NIC203 | chrV | 12015684 | 12016495 |
| sslp-4.3_NIC203 | FUN_019302 | Ctro | NIC203 | chrV | 12017693 | 12019061 |
| sslp-12_NIC203 | FUN_019312 | Ctro | NIC203 | chrV | 12046851 | 12047707 |
| sslp-5_NIC203 | FUN_019341 | Ctro | NIC203 | chrV | 12098633 | 12099142 |
| sslp-4.4_NIC203 | FUN_019342 | Ctro | NIC203 | chrV | 12099778 | 12100940 |
| sslp-13_NIC203 | FUN_019763 | Ctro | NIC203 | chrV | 13030575 | 13031658 |
| sslp-14_NIC203 | FUN_019764 | Ctro | NIC203 | chrV | 13031931 | 13032906 |
| sslp-6_NIC203 | FUN_019765 | Ctro | NIC203 | chrV | 13033424 | 13034904 |
| sslp-15_NIC203 | FUN_019767 | Ctro | NIC203 | chrV | 13038508 | 13039762 |
| sslp-7.2_NIC203 | FUN_019768 | Ctro | NIC203 | chrV | 13040253 | 13041212 |
| sif-1_NIC203 | NA | NA | NIC203 | chrV | 13042883 | 13043567 |
| sslp-8_NIC203 | FUN_019776 | Ctro | NIC203 | chrV | 13074045 | 13076050 |
| sslp-1.2_NIC203 | FUN_019778 | Ctro | NIC203 | chrV | 13082351 | 13084230 |
| sslp-9_NIC203 | FUN_019782 | Ctro | NIC203 | chrV | 13099770 | 13101571 |
| sslp-10.1_NIC203 | FUN_019787 | Ctro | NIC203 | chrV | 13109828 | 13111507 |
| sslp-10.2_NIC203 | FUN_019790 | Ctro | NIC203 | chrV | 13121881 | 13123495 |
| sslp-11.1_NIC203 | FUN_019793 | Ctro | NIC203 | chrV | 13133129 | 13134378 |
| sslp-11.2_NIC203 | FUN_019800 | Ctro | NIC203 | chrV | 13148827 | 13150466 |
| sslp-1.1_EG6180 | FUN_006541 | Ctro | EG6180 | chrII | 8459829 | 8463191 |
| sslp-2_EG6180 | FUN_016791 | Ctro | EG6180 | chrV | 6454491 | 6457748 |
| sslp-3_EG6180 | FUN_018741 | Ctro | EG6180 | chrV | 11977241 | 11978433 |
| sslp-4.1_EG6180 | FUN_018764 | Ctro | EG6180 | chrV | 12025019 | 12026080 |
| sslp-4.2_EG6180 | FUN_018765 | Ctro | EG6180 | chrV | 12026888 | 12027642 |
| sslp-5_EG6180 | FUN_018796 | Ctro | EG6180 | chrV | 12088806 | 12089315 |
| sslp-4.4_EG6180 | FUN_018797 | Ctro | EG6180 | chrV | 12090019 | 12090683 |
| sslp-16_EG6180 | FUN_018820 | Ctro | EG6180 | chrV | 12138630 | 12140281 |
| sslp-17.2_EG6180 | FUN_019198 | Ctro | EG6180 | chrV | 12999178 | 13001576 |
| sslp-17.1_EG6180 | FUN_019197 | Ctro | EG6180 | chrV | 12999249 | 13000010 |
| sslp-17.3_EG6180 | FUN_019199 | Ctro | EG6180 | chrV | 13001464 | 13003314 |
| sslp-18_EG6180 | FUN_019201 | Ctro | EG6180 | chrV | 13007179 | 13008307 |
| sslp-19_EG6180 | FUN_019202 | Ctro | EG6180 | chrV | 13009972 | 13010529 |
| sslp-20_EG6180 | FUN_019203 | Ctro | EG6180 | chrV | 13011557 | 13013581 |
| sslp-21_EG6180 | FUN_019204 | Ctro | EG6180 | chrV | 13016066 | 13017303 |
| sslp-6_EG6180 | FUN_019206 | Ctro | EG6180 | chrV | 13019110 | 13019906 |
| sslp-22_EG6180 | FUN_019207 | Ctro | EG6180 | chrV | 13020519 | 13021789 |
| sslp-23_EG6180 | FUN_019209 | Ctro | EG6180 | chrV | 13025607 | 13027937 |
| sslp-24_EG6180 | FUN_019210 | Ctro | EG6180 | chrV | 13029222 | 13029869 |
| sslp-7.1_EG6180 | FUN_019211 | Ctro | EG6180 | chrV | 13030631 | 13031353 |
| sslp-7.2_EG6180 | FUN_019212 | Ctro | EG6180 | chrV | 13034036 | 13035177 |
| sslp-25_EG6180 | FUN_019213 | Ctro | EG6180 | chrV | 13036610 | 13037514 |
| sslp-7.3_EG6180 | FUN_019214 | Ctro | EG6180 | chrV | 13039251 | 13040018 |
| sslp-8_EG6180 | FUN_019221 | Ctro | EG6180 | chrV | 13072044 | 13073202 |
| sslp-1.2_EG6180 | FUN_019223 | Ctro | EG6180 | chrV | 13082755 | 13083174 |
| sslp-9_EG6180 | FUN_019226 | Ctro | EG6180 | chrV | 13099586 | 13101320 |
| sslp-10.1_EG6180 | FUN_019231 | Ctro | EG6180 | chrV | 13109796 | 13110997 |
| sslp-11.1_EG6180 | FUN_019234 | Ctro | EG6180 | chrV | 13118762 | 13120009 |

**Table S4. AlphaFold 3 predictions of interactions between the SGYN-1 antidote and *C. tropicalis* SKR proteins.**

| Model | IPTM |
| --- | --- |
| sgyn_skr1_model_0 | 0.79 |
| sgyn_skr5_model_3 | 0.78 |
| sgyn_skr5_model_2 | 0.78 |
| sgyn_skr5_model_1 | 0.78 |
| sgyn_skr5_model_0 | 0.78 |
| sgyn_skr1_model_4 | 0.78 |
| sgyn_skr1_model_3 | 0.78 |
| sgyn_skr1_model_2 | 0.78 |
| sgyn_skr1_model_1 | 0.78 |
| sgyn_1_skr20_model_4 | 0.78 |
| sgyn_1_skr20_model_2 | 0.78 |
| sgyn_1_skr20_model_1 | 0.78 |
| sgyn_1_skr20_model_0 | 0.78 |
| sgyn_skr5_model_4 | 0.77 |
| sgyn_1_skr20_model_3 | 0.77 |
| sgyn_skr16_model_1 | 0.73 |
| sgyn_skr16_model_0 | 0.73 |
| sgyn_skr16_model_4 | 0.72 |
| sgyn_skr16_model_3 | 0.72 |
| sgyn_skr16_model_2 | 0.72 |
| sgyn_skr17_model_2 | 0.71 |
| sgyn_skr17_model_1 | 0.71 |
| sgyn_skr17_model_0 | 0.7 |
| sgyn_skr17_model_3 | 0.69 |
| sgyn_skr17_model_4 | 0.68 |
| sgyn_skr12_2_model_0 | 0.64 |
| sgyn_skr12_2_model_4 | 0.62 |
| sgyn_skr12_2_model_3 | 0.62 |
| sgyn_skr12_2_model_2 | 0.62 |
| sgyn_skr12_2_model_1 | 0.62 |
| sgyn_skr9_model_0 | 0.58 |
| sgyn_skr13_3_model_1 | 0.58 |
| sgyn_skr13_3_model_4 | 0.57 |
| sgyn_skr13_3_model_3 | 0.57 |
| sgyn_skr13_3_model_2 | 0.57 |
| sgyn_skr13_3_model_0 | 0.57 |
| sgyn_1_skr21_model_0 | 0.57 |
| sgyn_skr9_model_3 | 0.56 |
| sgyn_skr9_model_2 | 0.56 |
| sgyn_skr9_model_1 | 0.56 |
| sgyn_skr7_model_1 | 0.56 |
| sgyn_skr7_model_0 | 0.56 |
| sgyn_skr13_1_model_2 | 0.56 |
| sgyn_skr13_1_model_1 | 0.56 |
| sgyn_skr13_1_model_0 | 0.56 |
| sgyn_1_skr21_model_3 | 0.56 |
| sgyn_1_skr21_model_2 | 0.56 |
| sgyn_1_skr21_model_1 | 0.56 |
| sgyn_skr9_model_4 | 0.55 |
| sgyn_skr13_1_model_4 | 0.55 |
| sgyn_skr13_1_model_3 | 0.55 |
| sgyn_skr12_1_model_0 | 0.55 |
| sgyn_skr7_model_4 | 0.54 |
| sgyn_skr7_model_3 | 0.54 |
| sgyn_skr7_model_2 | 0.54 |
| sgyn_skr13_2_model_2 | 0.54 |
| sgyn_skr13_2_model_0 | 0.54 |
| sgyn_skr12_1_model_2 | 0.54 |
| sgyn_skr12_1_model_1 | 0.54 |
| sgyn_skr14_model_0 | 0.53 |
| sgyn_skr13_2_model_4 | 0.53 |
| sgyn_skr13_2_model_3 | 0.53 |
| sgyn_skr13_2_model_1 | 0.53 |
| sgyn_skr12_1_model_4 | 0.53 |
| sgyn_skr12_1_model_3 | 0.53 |
| sgyn_skr14_model_1 | 0.52 |
| sgyn_skr14_model_2 | 0.51 |
| sgyn_skr14_model_3 | 0.5 |
| sgyn_skr8_model_2 | 0.49 |
| sgyn_skr8_model_1 | 0.49 |
| sgyn_skr8_model_0 | 0.49 |
| sgyn_skr10_2_model_2 | 0.49 |
| sgyn_skr8_model_4 | 0.48 |
| sgyn_skr8_model_3 | 0.48 |
| sgyn_skr14_model_4 | 0.47 |
| sgyn_skr10_2_model_4 | 0.47 |
| sgyn_skr10_2_model_1 | 0.47 |
| sgyn_skr10_2_model_0 | 0.47 |
| sgyn_skr10_2_model_3 | 0.46 |
| sgyn_skr15_model_0 | 0.39 |
| sgyn_skr15_model_1 | 0.38 |
| sgyn_skr15_model_4 | 0.37 |
| sgyn_skr15_model_3 | 0.37 |
| sgyn_skr15_model_2 | 0.37 |
| sgyn_skr19_model_0 | 0.28 |
| sgyn_skr4_model_1 | 0.26 |
| sgyn_skr4_model_3 | 0.25 |
| sgyn_skr4_model_2 | 0.25 |
| sgyn_skr4_model_0 | 0.25 |
| sgyn_skr4_model_4 | 0.24 |
| sgyn_1_skr21_model_4 | 0.22 |
| sgyn_skr3_model_0 | 0.19 |
| sgyn_skr3_model_3 | 0.18 |
| sgyn_skr3_model_2 | 0.18 |
| sgyn_skr3_model_1 | 0.18 |
| sgyn_skr2_model_1 | 0.18 |
| sgyn_skr3_model_4 | 0.17 |
| sgyn_skr2_model_0 | 0.17 |
| sgyn_skr6_model_2 | 0.16 |
| sgyn_skr6_model_1 | 0.16 |
| sgyn_skr6_model_0 | 0.16 |
| sgyn_skr2_model_3 | 0.16 |
| sgyn_skr2_model_2 | 0.16 |
| sgyn_skr2_model_4 | 0.15 |
| sgyn_skr6_model_4 | 0.14 |
| sgyn_skr6_model_3 | 0.14 |
| sgyn_skr19_model_2 | 0.11 |
| sgyn_skr19_model_1 | 0.11 |
| sgyn_skr19_model_3 | 0.1 |
| sgyn_skr19_model_4 | 0.09 |

**Table S5. X-ray diffraction data collection and crystallographic refinement statistics for the THOR-1–SIF-1 complex.** Values in parentheses correspond to the highest-resolution shell.

| **Data collection** |  |
| --- | --- |
| Beamline | ESRF ID23-2 |
| Wavelength (Å) | 0.873 |
| Resolution range (Å) | 50.00 - 2.20<br>(2.33 - 2.20) |
| Space group | P 21 21 2 |
| Unit cell<br>a, b, c (Å)<br>$\alpha$, $\beta$, $\gamma$ (°) | 90.13, 98.48, 40.46<br>90, 90, 90 |
| Total reflections | 227,371 (22,685) |
| Unique reflections | 18,966 (2,982) |
| Multiplicity | 11.99 (7.61) |
| Completeness (%) | 99.36 (93.58) |
| Mean I/sigma(I) | 11.77 (1.02) |
| R-meas | 0.13 (1.31) |
| CC1/2 | 0.999 (0.547) |
| **Refinement** |  |
| Resolution range (Å) in refinement | 20-2.20 (2.26-2.20) |
| Reflections used in refinement | 18,843 (1,341) |
| Reflections used for R-free | 1,886 (134) |
| R-work | 0.187 |
| R-free | 0.231 |
| Number of heavy atoms in proteins | 2,357 |
| Number of heavy atoms in solvent | 192 |
| RMS (bonds) | 0.004 |
| RMS (angles) | 0.65 |
| Ramachandran favored (%) | 95.7 |
| allowed (%) | 4.3 |
| outliers (%) | 0 |

**Table S7. Pairs of *C. tropicalis* isolates carrying opposing toxin–antidote haplotypes collected at identical geographic coordinates.**

| isolate 1 | isolate 2 | region | coordinates | isolation date | substrate |
| --- | --- | --- | --- | --- | --- |
| NIC1915 | NIC1976 | Martinique | -61.0905,<br>14.6764 | 26.05.19 | Rotting_flower |
| NIC1939 | NIC1941 | Martinique | -61.1473,<br>14.8058 | 28.05.19 | Rotting_nut/pod/seed/frui<br>t |
| NIC747 | NIC748 | Dominica | -61.2969,<br>15.3157 | 14.05.14 | Rotting_nut/pod/seed/frui<br>t |
| NIC114 | NIC121 | Guadelou<br>pe | -61.6430,<br>16.0435 | 01.03.10 | Rotting_nut/pod/seed/frui<br>t |
| JU2750 | JU2752 | Guadelou<br>pe | -61.6951,<br>16.1761 | 09.06.14 | Rotting_nut/pod/seed/frui<br>t |

**Table S10. Summary of all primers and guide RNAs, and HDR donor templates.**

| Primers |  |  |  |
| --- | --- | --- | --- |
| Purpose | Primer 1 | Primer 2 | Product size |
| genotyping NIC203 vs NIL Chr. V; recombinant line screening (~13 Mb) | CCAACGAGTCAAGACGGTT<br>T | CAGTGACAAACACCCGAAAA | 373 bp<br>NIC203 /<br>276 bp<br>EG6180 |
| recombinant line screening (~12.7 Mb) | GCGTGTTGGCAAGTAGCAT<br>A | CCGACGGAAAACAAGAAAAA | 832 bp<br>NIC203 /<br>1064 bp<br>EG6180 |
| recombinant line screening (~13.2 Mb) | TCGTCGTAGGGCTCAAAAT<br>C | GTTTGCTGGGTGGCCTAGTA | 518 bp<br>NIC203 /<br>775 bp<br>EG6180 |
| *sif-1* mutants genotyping | GCCCTCAGAAAGGCTCTAC<br>GAA | GTGAGGAGTCGGGAAGTCAG<br>TC | 694 bp |
| *thor-1* deletion | GATTTTCCCGGTTTTCTGAC<br>GCT | AATCGGGGGAATAGGTTTCC<br>GG | 601 bp<br>del /<br>1970 bp<br>WT |
| *thor-1* frameshift genotyping | TGAATATGCCGACCCGTGA<br>GTT | ATTCTCAGCTGGTTCGAACGG<br>A | 363 bp |
| *loki-1* frameshift genotyping | ACGACTGTCTATCGCCTGC<br>TAC | TGGCGAGTTGTCTTCGTTGAC<br>TA | 590 bp |
| *sgyn-1* deletion genotyping | TGGCTATTTCACACGAAAAT<br>CGGA | AAAAGCCATTTCGCCACTACC<br>C | 463 bp<br>del /<br>1612 bp<br>WT |
| JU1373 vs EG6180 genotyping | GCCTAGAAAAACAATTGAT<br>GGCC | CGAGTAATTTCACCGGGTTG | 699 bp<br>JU1373 / |
|  |  |  | 556 bp<br>EG6180 |
| *sif-1* cloning from cDNA | ATGGAAGCTCCTCGTTGCA<br>ATGAAG | TTAAATGGCTTTGACGATCTC<br>GGCG | 552 bp |
| *thor-1* cloning from cDNA | ATGCCGACCCATTCTGCCC<br>ACCAG | TTATGACCTCGGACGAGCCAA<br>CATTAG | 405 bp |

| gRNAs and repair templates |  |  |  |
| --- | --- | --- | --- |
| Line | Modification | gRNA | Repair template |
| INK1263 | Arg<sup>142</sup>Glu; Arg<sup>151</sup>Glu; Thr<sup>147</sup>Ala / R142E; T147A; R151E | ACGGAAAGACGACA<br>TCCACA | GAAGAGGGCATCATAAAAGTCCGAGTC<br>GAACGAGAAGAGTATACTGAGACAGAT<br>GGTAAAGCTACTAGTACTGAGAAATGC<br>CACAACAGATACTTTCGAGAGGGACAA<br>GAGGAGATTTATG |
| INK1620 | Arg<sup>142</sup>Ala / R142A | GAAGAGTATACTAG<br>AACTGA | GGCATCATAAAAGTCCGAGTCGAACGA<br>GAAGAGTATACTGCTACTGACGGAAAG<br>ACGACATCCACAAGGAAATGCCACAAC<br>AGA |
| INK1551 | Arg<sup>151</sup>Ala / R151A | TCTGTTGTGGCATT<br>CCTTG | CGAGAAGAGTATACTAGAACTGACGGA<br>AAGACGACATCCACAGCTAAATGCCAC<br>AACAGATACTTTCGAGAGGGACAAGAG<br>GAGATT |
| INK1617 | Arg<sup>142</sup>Lys / R142K | GAAGAGTATACTAG<br>AACTGA | GGCATCATAAAAGTCCGAGTCGAACGA<br>GAAGAGTATACTAAGACTGACGGAAAG<br>ACGACATCCACAAGGAAATGCCACAAC<br>AGA |
| INK1545 | Arg<sup>151</sup>Lys / R151K | TCTGTTGTGGCATT<br>CCTTG | CGAGAAGAGTATACTAGAACTGACGGA<br>AAGACGACATCCACTAAAAAATGCCACA<br>ACAGATACTTTCGAGAGGGACAAGAGG<br>AGATT |
| INK1675 | Arg<sup>151</sup>Glu / R151E | TCTGTTGTGGCATT<br>CCTTG | GAGAAGAGTATACTAGAACTGACGGAA<br>AGACGACATCCACAGAAAAATGCCACA<br>ACAGATACTTTCGAGAGGGACAAGAGG<br>AGATT |
| INK1621 | Arg<sup>142</sup>Asn / R142N | GAAGAGTATACTAG<br>AACTGA | GGCATCATAAAAGTCCGAGTCGAACGA<br>GAAGAGTATACTAATACTGACGGAAAG<br>ACGACATCCACAAGGAAATGCCACAAC<br>AGA |
| INK1548 | Arg<sup>151</sup>Gln / R151Q | TCTGTTGTGGCATT<br>CCTTG | CGAGAAGAGTATACTAGAACTGACGGA<br>AAGACGACATCCACTCAAAAATGCCAC<br>AACAGATACTTTCGAGAGGGACAAGAG<br>GAGATT |
| INK463 | Arg<sup>142</sup>Glu / R142E | GAAGAGTATACTAG<br>AACTGA | GGCATCATAAAAGTCCGAGTCGAACGA<br>GAAGAGTATACTGAAACAGACGGAAAG<br>ACGACATCCACAAGGAAATGCCACAAC<br>AGATAC |
| INK231 | *thor-1* (-) | TGAAGCGACTCCAA<br>TCACAA | no |
| INK242 | *thor-1* (-); *sif-1* (-) | ATGATCACTCTTATG<br>ACATG | no |
| INK1310 | *thor-1* deletion | CATCACAAACCTTGT<br>TTCTCGT,<br>GCGTGGGAAAACCG<br>AATTTCT | CGCCGAAAATTGTCTTTTTGTTGAAAAA<br>GTTTGAAAAAAGGGTTTTCCATAATTTT<br>CCATCACAAACCTTTCCCACGCGAAAAT<br>TCATTTGAAAATCGAAAAGTAGCGCTGG<br>AGCGCGTTTGGAAGCTGCG |
| INK657 | *loki-1* (-) | ACGGATTGAGATTC<br>TGA | GGCTCCATTCGGCAAGTGACACTCTGC<br>CCGAAAAACCACGGATTGAGTAAGCTT<br>GACGGAAATTGATTTGATTGGAAGAGA<br>GGAGTTGATTGCAATCTTGG |
| INK1828 | *sgyn-1* (-) | GGCGCCAACTGATT<br>CTGAAAT,<br>GTCATTCCCTAGTAA<br>GTCTTC | GTCGTTTTGGGTAGTTTCTGCGGTCTC<br>GACTCTTGTTTCTTTGGGCGCCAACTAG<br>GGAATGACCAAAATGGGGTGTGAAAGT<br>TTGGTTCAAGACCTATACC |
